# Coordinated changes in glycosylation regulate the germinal centre through CD22

**DOI:** 10.1101/2021.04.30.442136

**Authors:** Jhon R. Enterina, Susmita Sarkar, Laura Streith, Jaesoo Jung, Britni M. Arlian, Sarah J. Meyer, Hiromu Takematsu, Changchun Xiao, Troy A. Baldwin, Lars Nitschke, Mark J. Shlomchick, James C. Paulson, Matthew S. Macauley

## Abstract

Germinal centres (GC) are sites of B-cell expansion and selection, which are essential for antibody affinity maturation. Compared to naive follicular B-cells, GC B-cells have several notable changes in their cell surface glycans. While these changes are routinely used to identify the GC, functional roles for these changes have yet to be ascribed. Detection of GCs by the antibody GL7 reflects a reduction in the glycan ligands for CD22, which is an inhibitory co-receptor of the B-cell receptor (BCR). To test a functional role for downregulated CD22 ligands in the GC, we generated a mouse model that maintains CD22 ligands on GC B-cells. With this model, we demonstrate that glycan remodeling is crucial for proper GC B-cell response, including plasma cell output and affinity maturation of antibodies. The defect we observe in this model is dependent on CD22, highlighting that coordinated downregulation of CD22 ligands on B cells plays a critical function in the GC. Collectively, our study uncovers a crucial role for glycan remodeling and CD22 in B-cell fitness in the GC.

## Introduction

Efficient clearance of invading pathogens partly relies on the production of high-affinity antibodies. These antibodies are secreted by plasma cells that emerge from microenvironments known as germinal centres (GCs), which transiently form in the secondary lymphoid organs following an infection or vaccination (*1*, *2*). GCs are anatomically organized into two functionally distinct compartments: the dark zone (DZ) and light zone (LZ). Within these compartments, GC B-cells undergo iterative clonal expansion, diversification of their B-cell receptor (BCR) genes through activation-induced cytidine deaminase (AID)-mediated somatic hypermutation (SHM) (*3*–*5*), and selection for memory B-cell and plasma cell differentiation (*5*–*7*). Positive selection relies on the ability of GC B-cells to successfully internalize, process, and present antigenic epitopes to cognate CD4^+^ follicular helper T-cells (T_FH_) for additional positive signals in the form of cytokines and co-stimulatory receptor interactions (*5*, *8*, *9*). Altogether, these highly orchestrated events in the GC ensure a preferential maintenance of B-cells with high-affinity BCRs; thereby securing the generation of high-quality antibodies for long lasting humoral immunity.

Earlier studies revealed that crosslinking of BCR on GC B-cells is inefficient in phosphorylating downstream signaling components compared to their naive B-cell counterparts (*10*, *11*). Dampening of BCR signaling in GC B-cells is mediated by Src homology 2 (SH2) domain-containing protein-tyrosine phosphatase 1 (SHP1) (*12–14*). SHP1 negatively regulates BCR activation by binding to phosphorylated immunoreceptor tyrosine-based inhibitory motifs (ITIMs) of BCR inhibitory co-receptors, such as CD22, CD72, PIR-B, Siglec-10/G, FcγRIIb, and FCRL5 (*15*–*19*). Indeed, deletion of *Ptpn6*, the gene encoding SHP1 in mice, results in substantially reduced GC B-cell numbers and plasma cell output, which was largely due to increased apoptotic cell death (*10*, *20*, *21*). Overall, these data strongly suggest that effective inhibition of BCR signaling following antigen engagement is essential for a proper GC response. However, it remains unclear as to how BCR signal strength is differentially regulated between naive and GC B-cells and whether unique changes on upstream receptors of SHP1 exist in the GC, resulting in recruitment of this key phosphatase to the BCR.

In the GC, antigen-activated B-cells display a dramatic remodeling of glycans on cell surface glycoproteins. In particular, several defining events relate to changes in the monosaccharide sialic acid, which caps cell surface N- and O-linked glycans. These changes are so distinct that two of the most widely used agents to identify mouse GC B-cells recognize and bind to glycan structures. One is the plant lectin peanut agglutinin (PNA) that binds to B-cells in the GC due to the downregulation of sialyltransferase ST3Gal1, resulting in appearance of its ‘unsialylated ligand’ (Galβ1-3GalNAcαThr/Ser) on O-linked glycans (*22*). Second is the glycan epitope recognized by the antibody GL7, which emerges from the downregulation of the enzyme CMP sialic acid hydroxylase (CMAH) in GC B-cells (*23*). GL7 detects glycans terminating in α2-6-linked *N*-acetylneuraminic acid (Neu5Ac) (*23*, *24*). Downregulation of CMAH prevents the hydroxylation of cytidine monophosphate (CMP)-Neu5Ac and its conversion to CMP-*N*-glycolylneuraminic acid (CMP-Neu5Gc), which is preferentially expressed on naive murine B-cells. The switch from Neu5Gc to Neu5Ac on GC B-cells also has a profound effect on ligands of mouse CD22 (mCD22). Specifically, mCD22 binds to α2-6-linked Neu5Ac-containing glycans 20-fold weaker than their corresponding α2-6-linked NeuGc-containing counterparts (*24*).

The impact of the loss of ligand recognition by mCD22 on naive B-cells can be seen in studies examining B-cell responses from mice that have disruptions in the interaction of CD22 with its ligands. Specifically, deletion of the sialyltransferase ST6Gal1 or knock-in of a mutant CD22 (R130E) that can’t recognize its glycan ligands leads to blunted B-cell responses in the naive B-cell compartment (*25*, *26*). Super resolution imaging studies have supported these findings, demonstrating that CD22-glycan interactions keep CD22 within nanoclusters, which also contain CD45 and galectin-9, and away from the BCR (*27, 28*). As consequence, ablating CD22-ligand interactions results in increased CD22-BCR association and dampened BCR signaling. Based on these established roles for CD22-glycan interactions, we hypothesized that downregulation of CD22 glycan ligands on GC B-cells likewise plays a similar role, which could serve to increase the ability of CD22 to antagonize BCR signaling.

Interestingly, humans have lost the ability to biosynthesize Neu5Gc due to an exon deletion in the *CMAH* gene (*29, 30*). However, an alternative mechanism leads to downregulation of CD22 ligands on human GC B-cells. Human CD22 has evolved to recognize a sulfated form of its glycan ligand (Neu5Acα2–6Galβ1–4(6-sulfo)GlcNAc) and several lines of evidence have shown that this glycan structure shows substantial staining on naive and memory B-cells but not on GC B-cells (*24*, *31*). Thus, downregulation of CD22 ligands in the GC is an evolutionarily conserved mechanism catering to the ligand specificity of CD22 from mouse and human. Given this evolutionary conserved process, we hypothesized that modulation of CD22 ligands in GC B-cells plays an important role. To test this, we developed a transgenic mouse model that conditionally expresses CMAH in B-cells, thereby preventing downregulation of CD22 ligands. Here, we show that constitutive expression of CMAH impairs the GC response, which is CD22-dependent, and that a similar defect is observed in CD22^KO^ mice. The absence of CD22 or inability to downregulate CD22 ligands leads to higher BCR activation, defective antibody affinity maturation, and increased apoptotic cell death in GC B-cells. Collectively, our findings strongly suggest that glycan remodeling, through downregulation of CMAH in the GC, is crucial for the maintenance and competitiveness of GC B-cells, underscoring GC-specific downregulation of CD22 ligands as a critical mechanism contributing to rewiring of BCR signaling in the GC.

## Results

### Constitutive expression of CMAH maintains high affinity ligands for CD22 on GC B-cells

Appearance of the GL7 glycan epitope is mediated by loss of the enzyme CMAH on *in vitro* stimulated primary murine B-cells (*23*). To confirm whether appearance of the GL7 epitope on GC B-cells occurs through transcriptional downregulation of *Cmah*, we compared the mRNA expression levels of *Cmah* in mouse DZ and LZ GC B-cells to follicular (non-GC) B-cells using a publicly available dataset (GSE109125; (*32*)). Follicular B-cell levels of *Cmah* transcript are indeed significantly downregulated in both DZ and LZ GC B-cells (**Fig. 1A**). To explore the functional role for this switch in sialic acid from Neu5Gc to Neu5Ac as naive B-cells differentiate into GC B-cells, we developed a transgenic mouse model to modulate Neu5Gc expression in a cell-type specific manner, for the purpose of preventing downregulation of Neu5Gc in GC B-cells. *Cmah* was knocked into the *Rosa26* locus with an upstream loxP-STOP-loxP cassette to enable Cre recombinase-dependent expression of CMAH, which we describe as *R26^lsl^-Cmah* mice (**Fig. 1B**). To characterize the effect of constitutive CMAH expression on glycans expressed by GC B-cells, we initially crossed *R26^lsl^-Cmah* mice with *Mb1^Cre^* mice (*33*) to drive constitutive CMAH expression in the B-cell lineage. As anticipated, immunization of *Mb1^Cre^*×*R26^lsl^-Cmah*, denoted hereafter as CMAH^ON^, mice with a T-dependent antigen resulted in GC B-cells (B220^+^CD19^+^IgD^−^CD38^low^Fas^+^PNA^+^ cells) that do not express the GL7 epitope (**Fig. 1C, D**) and maintain Neu5Gc expression at comparable levels to naive B-cells (B220^+^CD19^+^IgD^+^ cells) (**Fig. 1E, F**). Maintaining CMAH expression in GC B-cells also prevented the downregulation of CD22 ligands, as demonstrated by staining with murine CD22-Fc (**Fig. 1G, H**).

**Fig 1.**
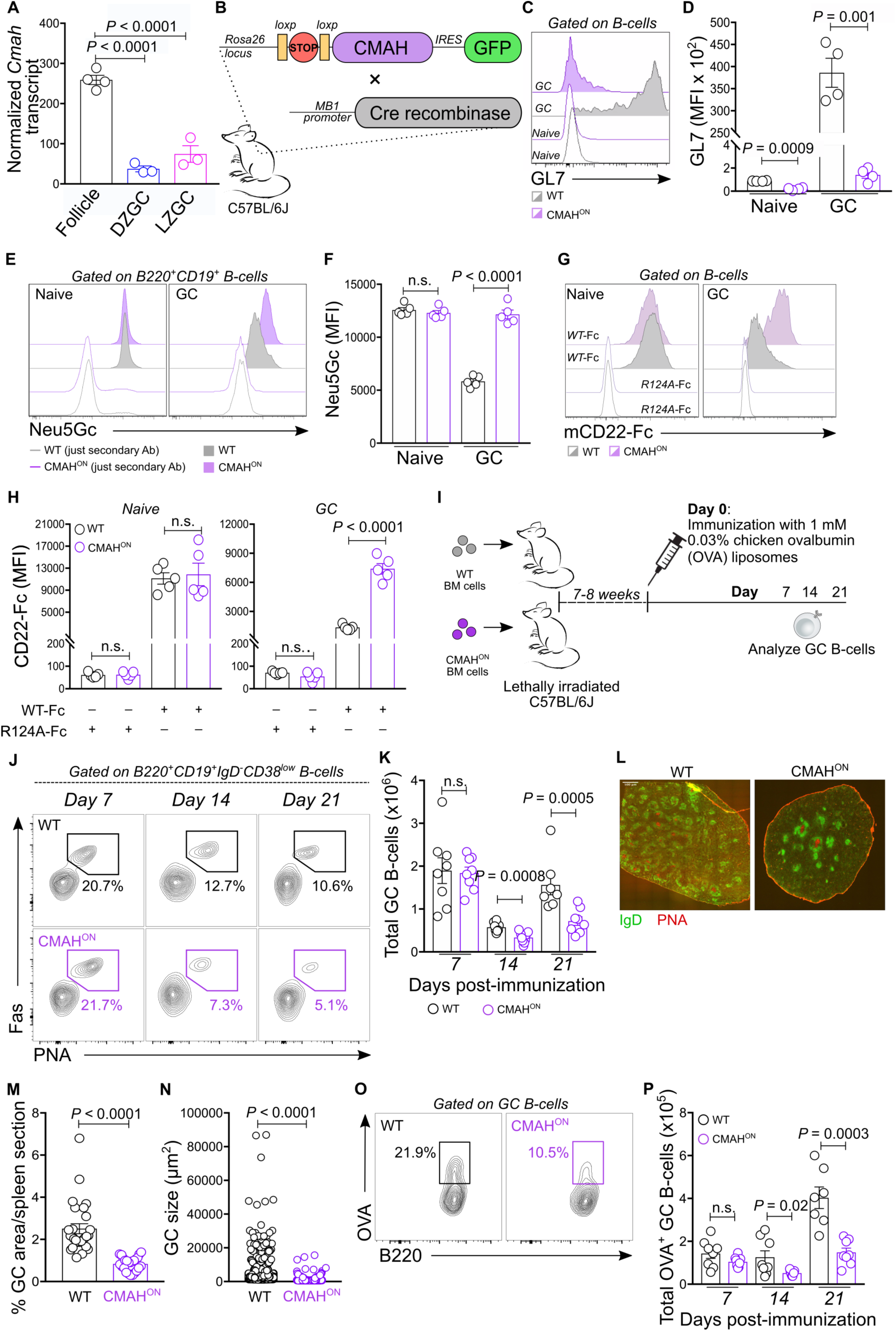
Constitutive expression of CMAH in GC B-cells restricts GC B-cell response following immunization with T-dependent antigen. (**A**). *Cmah* gene transcript is significantly downregulated in mouse dark zone (DZ) and light zone (LZ) GC B-cells compared to follicular B-cells (source: GSE109125). (**B**) Graphical representation of a *Cmah* transgenic mouse model developed for this study. Cre recombinase-inducible CMAH transgene was inserted in the *Rosa26* locus and expresses green fluorescent protein (GFP) under the control of an internal ribosome entry site (IRES). (**C,D**) Flow cytometric analysis of GL7 epitope expression on WT and CMAH transgene-expressing naive and GC B-cells (C) and quantification of GL7 epitope levels (D). (**E,F**) Flow cytometric analysis of Neu5Gc sialic acid levels on WT and CMAH transgene-expressing (*Mb1Cre*×*R26lsl-Cmah* or CMAHON) naive and GC B-cells (E) and quantification of the Neu5Gc levels (F). (**G,H**) Flow cytometric analysis of mouse CD22-Fc or CD22 R124A-Fc staining on WT and CMAHON naive and GC B-cells (G) and quantification of mCD22-Fc or CD22 R124A-Fc staining on WT and CMAHON naive and GC B-cells (H). (**I**) Immunization scheme of *R26lsl*-*Cmah* (WT) and CMAHON chimera mice. Following immunization of T-dependent antigen, spleens were collected at various time points to analyze for GC B-cells. (**J**) Flow cytometric gating strategy for total GC B-cells. (**K**) Quantification of the absolute number of *R26lsl-Cmah* (WT) and CMAHON GC B-cells in immunized mouse spleens. (**L**) Immunofluorescence images of spleen section from immunized WT and CMAHON mice, at day 14 post-immunization. (**M,N**) Quantification of percent GC area per spleen section (M) and GC cluster sizes (N) in spleens of immunized WT or CMAHON mice. (**O**) Gating strategy for identification of OVA-specific GC B-cells in immunized *R26lsl*-*Cmah* (WT) and CMAHON mice spleens. (**P**) Quantification of the absolute number of OVA+ *R26lsl*-*Cmah* (WT) and CMAHON GC B-cells in spleens of immunized mice.

### CMAH expression restricts B-cell responses in the GC

To test the impact of maintaining CMAH expression in GC B-cells, lethally irradiated WT B6 mice were reconstituted with bone marrow (BM) cells from either *R26^lsl^-Cmah* (as WT control) or CMAH^ON^ mice to generate chimera mice, to control the age and sex of the mice for large cohort. Reconstituted mice were immunized with liposomes displaying chicken ovalbumin (OVA) antigen (**Fig. 1I**). This immunogen gives a robust and consistent T-dependent antibody response, while avoiding the inclusion of a TLR agonist (*34*–*36*). Flow cytometric analysis revealed that the total number of GC B-cells in immunized CMAH^ON^ mice is comparable to their WT counterparts on day 7 post-immunization (**Fig. 1J, K**). However, on days 14 and 21 after immunization, we observed a significant reduction in the number of GC B-cells from immunized CMAH^ON^ mice (**Fig. 1K**). Immunofluorescence analysis of spleens, collected 14 days post-immunization, identified a decrease in the size of GC clusters (IgD^−^PNA^+^) in CMAH^ON^ mice compared with WT mice, validating the phenotype we observed using flow cytometry (**Fig. 1L-M**). Concurrently, we assessed the generation of OVA-specific GC B-cells after immunization. Similar to what we observed with total GC B-cells, we found normal levels of OVA^+^ GC B-cells between WT and CMAH^ON^ on day 7, but at later time points, CMAH^ON^ mice had fewer OVA^+^ GC B-cells than their WT counterparts (**Fig. 1O,P**). Within a mixed BM chimera, constitutive expression of CMAH in the B-cells led to an obvious competitive disadvantage in the GC, which was noticeable even at one week post-immunization (**Fig. 2A, B**). Overall, these findings suggest loss of CMAH expression, and consequently Neu5Gc, is critical for a proper B-cell response in the GC.

**Fig 2.**
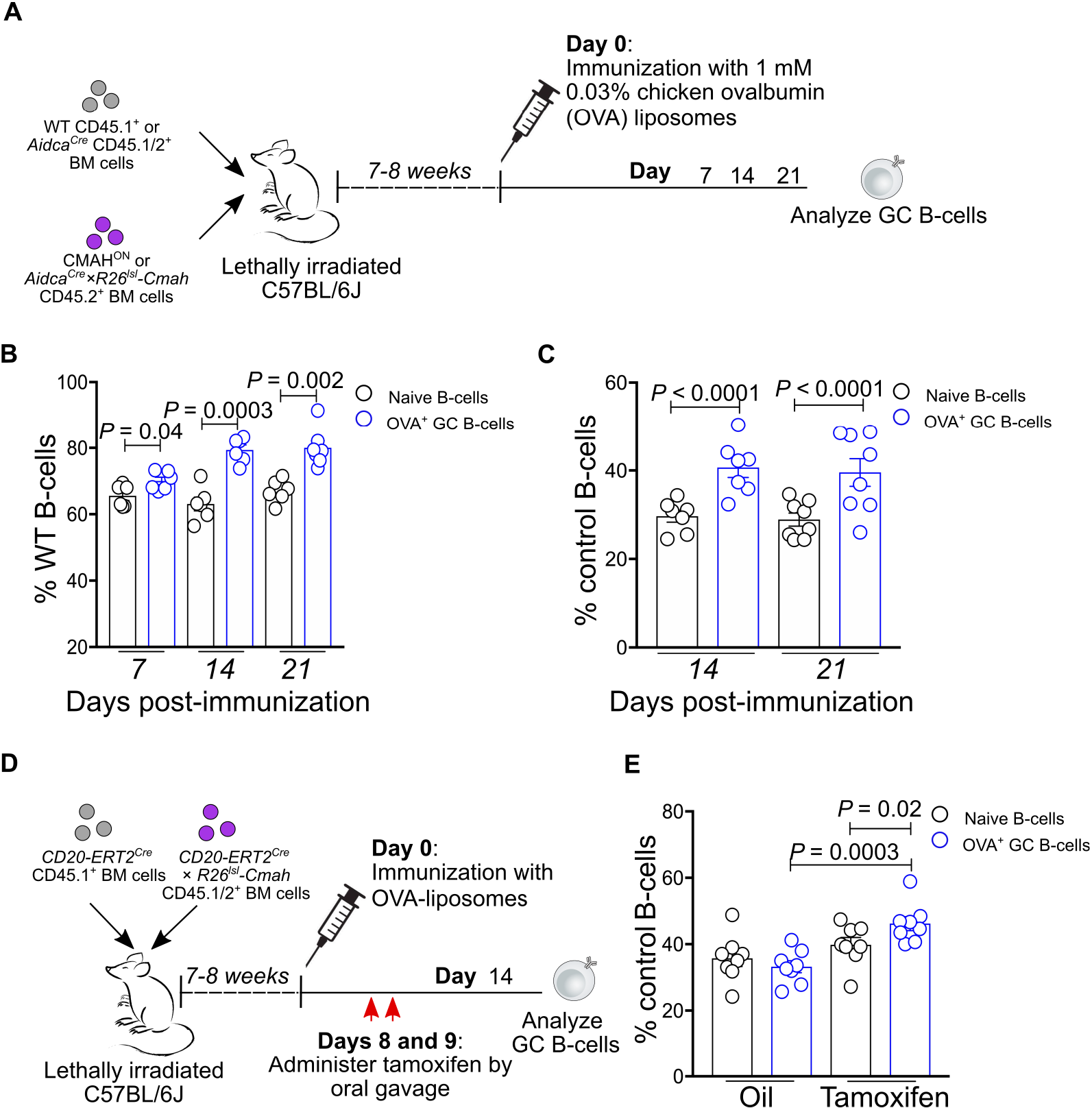
Impact of CMAH in restricting GC B-cell responses is GC specific. (**A**) Scheme for mixed bone marrow generation and immunization. (**B**) Quantification of percent WT naive and OVA+ GC B-cells in mixed bone marrow chimera mice reconstituted with 1:1 CD45.1+ WT and CMAHON BM cells. Spleens from immunized chimera mice were collected and analyzed by flow cytometry on days 7, 14, and 21 post-immunization. (**C**) Quantification of percent *AidcaCre* (control) naive and OVA+ GC in mixed bone marrow chimera mice reconstituted with 1:1 *AidcaCre* and *AidcaCre*×*R26lsl-Cmah* bone marrow cells. Spleens from immunized chimeras were collected and analyzed by flow cytometry on days 14 and 21 post-immunization. (**D**) Scheme for generating a tamoxifen-induced mixed chimera model for temporal expression of CMAH in GC B-cells. Equal number of *CD20-ERT2Cre* CD45.1+ and *CD20-ERT2Cre*×*R26lsl-Cmah* CD45.1/2+ bone marrow cells were transplanted in irradiated C57BL/6J mice. Reconstituted chimera mice were immunized with OVA-displaying liposomes and spleens were collected and analyzed by flow cytometry on day 14 post-immunization (day 5 after second tamoxifen or oil oral gavage). (**E**) Quantification of percent *CD20-ERT2Cre* CD45.1+ (control) naive and OVA+ GC B-cells in immunized mixed bone marrow chimera mice administered with oil (vehicle) and tamoxifen.

Given that CMAH^ON^ mice constitutively express CMAH in B-cells starting early in B-cell development, we aimed to test if this is a GC-specific phenotype by crossing *R26^lsl^-Cmah* mice with *Aidca^Cre^* mice to induce Cre expression 2-3 days post-immunization (*37*). Chimera mice were generated by transplanting *Aidca^Cre^* (WT control) and *Aidca^Cre^*×*R26^lsl^-Cmah* BM cells. After immunization, we found that OVA^+^ WT GC B-cells were outcompeting *Aidca^Cre^×R26^lsl^-Cmah* GC B-cells two weeks post-immunization (**Fig. 2C**). When expression of CMAH was temporally induced in the B-cell lineage by crossing human *CD20-ERT2^Cre^* and *R26^lsl^-Cmah* mice and administering tamoxifen on days 8 and 9 post-immunization (*12*), a competitive disadvantage of *CD20-ERT2^Cre^* over *CD20-ERT2^Cre^*×*R26^lsl^-Cmah* was also observed in the GC compartment (**Fig. 2D, E**). These findings demonstrate that downregulation of CMAH expression is essential for the maintenance of B-cells in the GC.

### Impaired GC response in CMAH transgenic B-cells is dependent on CD22

The function of mCD22 is regulated by the presence of its *cis*-ligands, and the preferred glycan ligands of CD22 require CMAH (*24*). Therefore, we tested whether sustained expression of CMAH in GC B-cells negatively impacts the GC response through CD22 by crossing CD22^KO^ mice and CMAH^ON^ mice. Mixed BM chimeras were set up using cells from CD22^KO^ and CD22^KO^×CMAH^ON^ donor mice. Following immunization, the previously observed competitive disadvantage in CMAH^ON^ GC B-cells was no longer observed (**Fig. 3A**). To test this in an adoptive transfer model, we used hen egg lysozyme (HEL)-specific B-cells on a CD22^KO^ and CD22^KO^×CMAH^ON^ background and tracked the formation of HEL^+^ GC B-cells in competition (**Fig. 3B**). After immunization with HEL-displaying liposomes, the ratio of adoptively transferred cells was comparable with their unimmunized counterparts (**Fig. 3C**). In contrast, immunization of mice adoptively transferred with HEL-specific B-cells on a WT and CMAH^ON^ background displayed a clear competitive disadvantage for CMAH^ON^ GC B-cells (**Fig. 3D**). Together, these experiments support the concept that the defective phenotype observed in CMAH-expressing GC B-cells is dependent on CD22.

**Fig 3.**
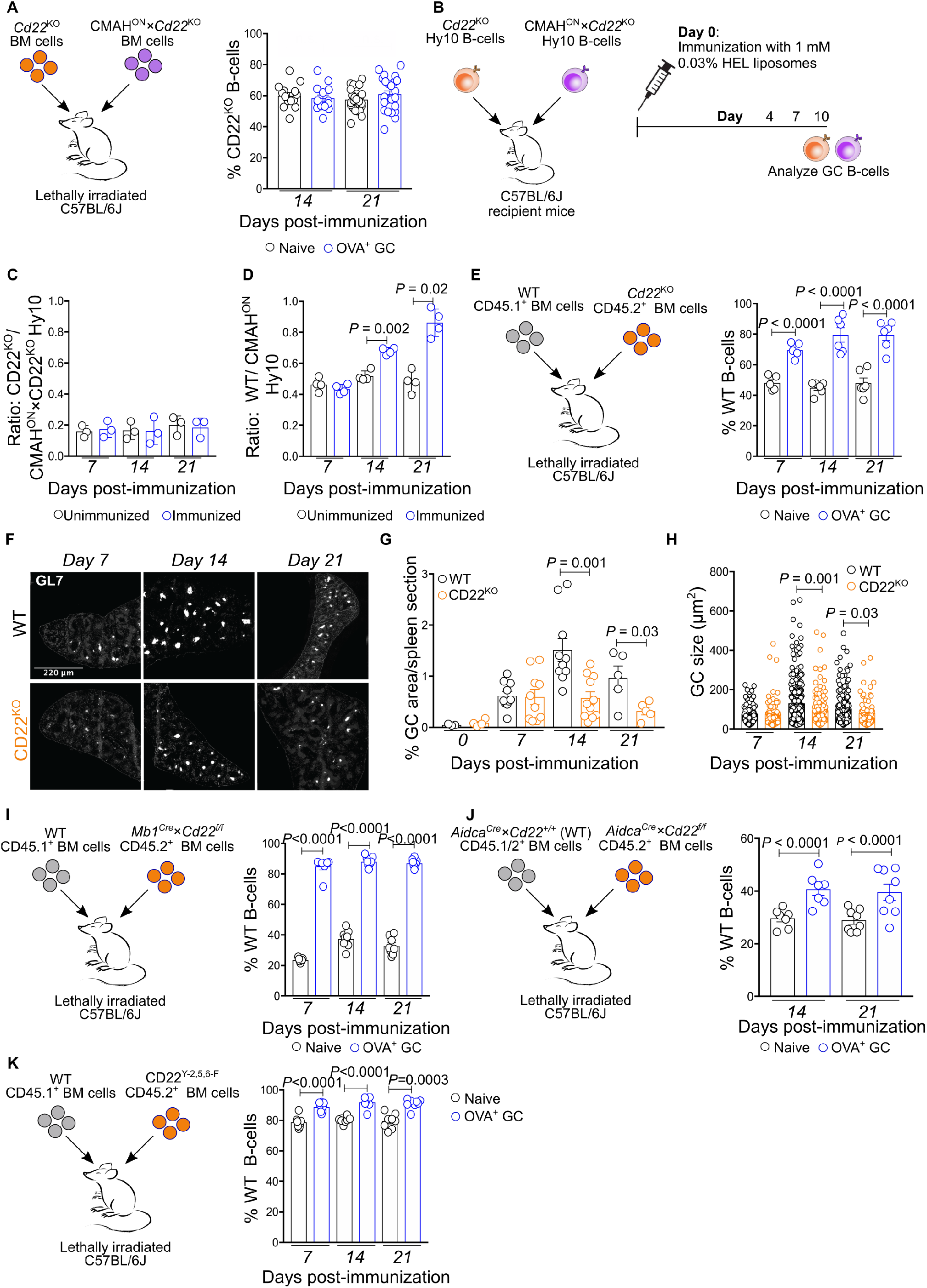
Restricted B-cell response in CMAH-expressing GC B-cells is dependent on CD22. (**A**) Quantification of percent CD22KO GC B-cells in mixed bone marrow chimera mice reconstituted with BM cells from *Cd22*KO and CMAHON×*Cd22*KO donor mice, after days 14 and 21 post-immunization. (**B**) Scheme for the adoptive transfer of HEL-specific B-cells into recipient mice and immunization of these mice with HEL antigen displayed on liposomes. (**C**) Quantification of percent HEL-specific *Cd22*KO B-cells in unimmunized and immunized mice that have been adoptively transferred with HEL-specific CD22KO and CMAHON×*Cd22*KO B-cells. (**D**) Quantification of percent HEL-specific WT B-cells in unimmunized and immunized mice that have been adoptively transferred with HEL-specific WT and CMAHON B-cells. (**E**) Quantification of percent WT GC B-cells in mixed bone marrow chimera mice, reconstituted with BM cells from CD45.1+ WT and CD22KO donor mice, after days 7, 14, and 21 post-immunization. (**F-H**) Immunofluorescence imaging of GCs (GL7+) (**F**) and quantifications of percent area covered by GC clusters/section (**G**) and size of GC clusters (**H**) in spleens of immunized WT and CD22KO mice. (**I**) Quantification of percent WT GC B-cells in mixed bone marrow chimera mice, reconstituted with BM cells from CD45.1+ WT and CMAHON donor mice, after days 7, 14, and 21 post-immunization. (**J**) Quantification of percent WT GC B-cells in mixed bone marrow chimera mice, reconstituted with BM cells from *AidcaCre*×*Cd22+/+* (as WT control) and *AidcaCre*×*Cd22f/f* donor mice, after days 7, 14, and 21 post-immunization. (**K**) Quantification of percent WT GC B-cells in mixed bone marrow chimera mice, reconstituted with BM cells from CD45.1+ WT and CD22Y-2,5,6-F donor mice, after days 7, 14, and 21 post-immunization.

### CD22 is critical for B-cell competitiveness in the GC

To test whether CD22 is critical for B-cell responses in the GC under more physiologically-relevant conditions (in the absence of constitutive CMAH expression), we generated mixed chimera mice transplanted with WT and *Cd22*^KO^ BM cells and then immunized them with OVA-displaying liposomes. OVA^+^ WT GC B-cells significantly outcompeted OVA^+^ CD22^KO^ GC B-cells even starting on day 7 post-immunization (**Fig. 3E**). Immunofluorescence imaging studies of spleens from immunized WT and CD22^KO^ mice, at varied time points post-immunization, also revealed the significantly reduced GC size (GL7^+^) in CD22^KO^ mice compared with their WT counterparts (**Fig. 3F-H**), confirming our flow cytometry data. To assess whether this defective GC B-cell response observed with CD22^KO^ B-cells is GC-specific, we generated floxed CD22 (*Cd22^f/f^*) mice and crossed them first with *Mb1^Cre^* mice. Following immunization of mixed chimera mice transplanted with WT and *Mb1^Cre^*×*Cd22^f/f^* BM cells, antigen-specific WT B-cells outcompeted the *Mb1^Cre^*×*Cd22^f/f^* B-cells in the GC, validating the GC defect observed in CD22^KO^ mice is B-cell intrinsic (**Fig. 3I**). We next crossed *Cd22^f/f^* mice with *Aidca^Cre^* mice. Tracking antigen-specific GC B-cell formation in immunized mixed chimera mice reconstituted with *Aidca^Cre^* (WT control) and *Aidca^Cre^*×*Cd22^f/f^* BM cells, we observed a competitive disadvantage for *Aidca^Cre^*×*Cd22^f/f^* GC B-cells on days 14 and 21 post-immunization (**Fig. 3J**). To assess whether the impaired GC response in CD22^KO^ B-cells is dependent on its ITIMs, we tracked the antigen-specific GC B-cell response of WT and CD22^Y-2,5,6-F^ B-cells, which have knock-in mutations in three of its key cytoplasmic ITIMs (*26*). Following immunization, we identified that loss of functional ITIMs on CD22 resulted in poor GC B-cell response, recapitulating a similar phenotype seen in CD22^KO^ GC B-cells (**Fig. 3K**). Collectively, these results strongly suggest that regulation of signaling by CD22 is required for the maintenance of B-cells in the GC.

### Loss of CD22 impairs interzonal distribution of GC B-cells

Repeated migration of GC B-cells between the DZ and LZ ensures the maintenance and selection of clones with high affinity BCRs. To test whether CMAH expression or loss of CD22 affects DZ/LZ transition of GC B-cells, we probed DZ (CXCR4^high^CD86^low^) and LZ (CXCR4^low^CD86^high^) GC B-cells using flow cytometry (**Fig. S1A**). After immunization, we found a normal distribution of CMAH^ON^ GC B-cells in the DZ and LZ compartments (**Fig. S1B**). However, CD22^KO^ GC B-cells displayed less LZ GC B-cells than WT GC B-cells (**Fig. S1C**). Consistent with the result by flow cytometry, immunofluorescence staining of spleens showed that the % of GC B-cells in the LZ (IgD^−^PNA^+^CD21/35^+^), compared to the total GC, was significantly lower in CD22^KO^ mice when compared with WT and CMAH^ON^ mice (**Fig. S1D,E**).

### Ablation of CD22 and constitutive expression of CMAH impairs antibody affinity maturation

To evaluate whether the defective GC B-cell response observed in CD22^KO^ and CMAH transgenic models results in defective humoral immunity, we immunized WT, CD22^KO^, and CMAH^ON^ mice with liposomes that display OVA conjugated with NP-hapten (**Fig. 4A**). Antibody affinity maturation was determined by monitoring high affinity anti-NP IgG1 and total anti-NP IgG1, using NP_2_-BSA and NP_23_-BSA, respectively, by ELISA at different time points after immunization. The NP_2_:NP_23_ ratio of the *IC*_50_ values in the ELISAs were used to estimate antibody affinity maturation, with a higher NP_2_:NP_23_ ratio being indicative of better antibody affinity maturation. We found that deficiency for CD22 impaired the maturation of anti-NP antibodies and were not able to recover even at a later time point (day 42) post-immunization. CMAH^ON^ mice also displayed a significant, yet modest, reduction in antibody affinity maturation at earlier time points post-immunization, but this difference compared to WT mice was no longer apparent by day 42 (**Fig. 4B**). To investigate whether this defective production of high affinity antibodies in CD22^KO^ mice is due to diminished SHM, NP-specific GC B-cells (B220^+^CD19^+^IgD^−^ GL7^+^NP_20_+ cells) were isolated by FACS from WT and CD22^KO^ mice, at 21 days post-immunization with NP-OVA displaying liposomes. The V_H_186.2 region was sequenced and found to have a similar mutation rate between WT and CD22^KO^ (**Fig. 4C-E**). Additionally, we also observed a comparable number of clones between WT (*n*=15/33) and CD22^KO^ (*n*=17/32) with the affinity enhancing mutations, W33L and K59R (*38*) (**Fig. 4F**).

**Fig 4.**
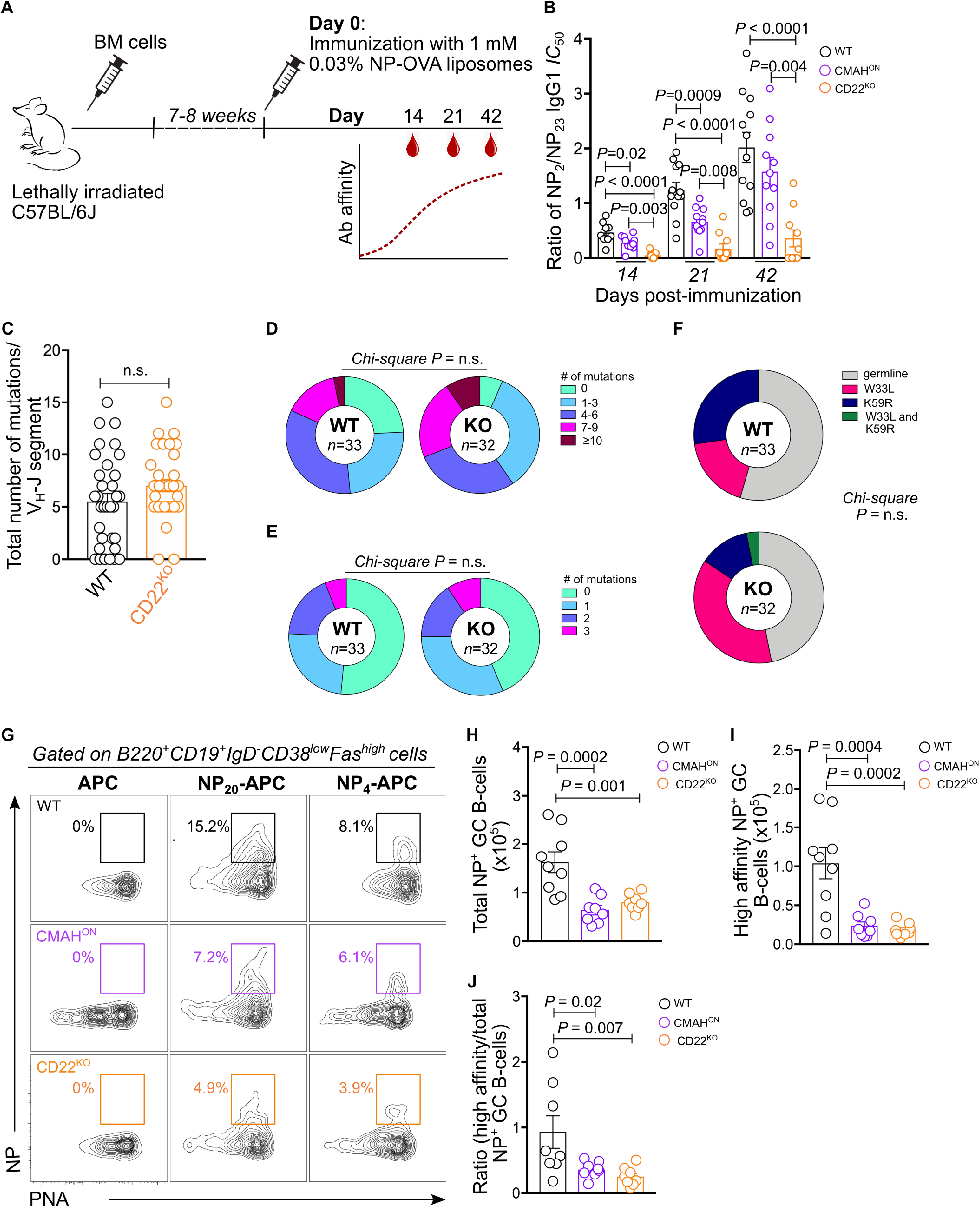
CD22 is not required in BCR somatic hypermutation but crucial for selection or maintenance of high-affinity antigen-specific GC B-cell clones as well as antibody affinity maturation. (**A**) Schematic of the immunization procedure used to assess antibody affinity maturation. (**B**) Estimation of affinity maturation of NP-specific IgG1 from sera of immunized WT, CMAHON, and CD22KO mice collected at different time points post-immunization (**C**) Quantification of total number of mutations in the V_H_-J segment of NP^+^ WT and CD22^KO^ GC B-cells. (**D**) Distribution of non-silent mutations present in the V region of NP+ WT and CD22KO GC B-cells. (**E**) Distribution of non-silent mutations present in the Junction region of NP+ WT and CD22KO GC B-cells. (**F**) Distribution of affinity enhancing mutations present in the V region of NP+ WT and CD22KO GC B-cells. (**G**) Flow cytometric gating strategy used to determine the numbers of high-affinity NP+ and total NP+ GC B-cells in spleens of immunized WT, CMAHON, and CD22KO mice. (**H,I**) Absolute number of total NP+ (H) and high affinity NP+ (I) GC B-cell present in spleens of immunized WT, CMAHON, and CD22KO mice. (**J**) Ratio of high-affinity over total NP+ GC B-cells in immunized WT, CMAHON, and CD22KO mice. *Chi*-square probability test was performed to test for statistical difference in mutation levels between WT and CD22KO (D-F).

To assess whether CD22 and its *cis*-ligands control the maintenance of GC B-cell clones with high affinity BCRs, we quantified the number of high affinity NP^+^ and total NP^+^ GC B-cells in WT, CD22^KO^, and CMAH^ON^ mice, immunized with NP-OVA displaying liposomes, by flow cytometry at day 14 post-immunization. We found that the absolute numbers of both high affinity NP^+^ and total NP^+^ GC B-cells, identified with NP_4_-APC and NP_20_-APC, respectively (**Fig. 4G**), were significantly lower in CD22^KO^ and CMAH^ON^ mice compared to WT (**Fig. 4H,I**). Likewise, the ratio of high affinity over total NP^+^ GC B-cells was also substantially reduced in CD22^KO^ and CMAH^ON^ mice (**Fig. 4J**). These findings demonstrate that CD22 and loss of its *cis*-ligands on GC B-cells are pivotal for the maintenance of B-cell clones with high affinity BCRs.

### Attenuated memory B-cell and plasma cell output in CD22^KO^ and CMAH^ON^ mice

Given that both CMAH^ON^ and CD22^KO^ mice show reduced numbers of antigen-specific GC B-cells following immunization, we investigated whether this finding correlates with decreased memory B-cell and plasma cell output as well. To test this, we quantified the number of NP^+^ memory B-cells (B220^+^CD19^+^IgD^−^ CD38^high^PNA^−^NP20+ cells) and plasma cells (CD138^high^CD19^−^NP_20_+ cells) using flow cytometry (**Fig. S2A**). We found that spleens of CMAH^ON^ and CD22^KO^ mice, after day 14 post-immunization with NP-OVA displaying liposomes, displayed remarkably lower numbers of both memory B-cells and plasma cells than their WT counterparts (**Fig. S2B-G**). These data indicate that decreased antibody responses observed in CMAH^ON^ and CD22^KO^ mice are likely due to reduced plasma cells emerging from the GC.

### Transcriptome analysis reveals a crucial role of CD22 in maintaining B-cell fitness in the GC

To determine the putative mechanisms responsible for the defective GC response in CD22^KO^ B-cells, we analyzed the impact of CD22 deficiency on the transcriptome profile of GC B-cells by RNA sequencing. The DZ (B220^+^CD19^+^IgD^−^GL7^+^CXCR4^high^CD86^low^) and LZ (B220^+^CD19^+^IgD^−^GL7^+^CXCR4^low^CD86^high^) GC B-cells from WT and CD22^KO^ mice, at day 14 post-immunization with OVA-displaying liposomes, were sorted using FACS and RNA was isolated and processed for next generation sequencing (**Fig. S3**). We found numerous differentially expressed genes (DEGs) that are either upregulated or downregulated in CD22^KO^ GC B-cells (**Fig. 5A-C**). Interestingly, we observed more DEGs in the LZ than in the DZ GC compartment. These include genes encoding for GIMAP GTPase family (e.g. *Gimap4, Gimap6,* and *Gimap7*; down), mitochondrial enzymes (e.g. *mt-Nd3, Ndufa1,* and *mt-Nd4l*; down), cell cycle regulatory proteins (e.g. *Ccnb1*, *Ccnb2*, and *Cdk6*; down), GC-relevant transcription factors (e.g. *Hhex, Tox2,* and *Irf1*; down), and B-cell activation signaling proteins (e.g. *Nfat5, Nr4a1,* and *Nfkbie*; up) (**Table S1**). To interrogate the distinct biological processes and pathways disrupted in the GC due to the loss of CD22, a gene enrichment analysis was performed using EnrichR with DEGs containing a *P* value < 0.05. We found that both DZ and LZ CD22^KO^ GC B-cells displayed defective oxidative phosphorylation, cell cycle progression, and cholesterol biosynthesis. On the other hand, upregulated genes in CD22^KO^ GC B-cells converged on pathways for B-cell activation, Toll-like receptor signaling, and apoptosis (**Fig. 5D, E**). Taken together, transcriptome analysis uncovers that expression of CD22 controls B-cell activation and other biological processes relevant for B-cell competitiveness in the GC.

**Fig 5.**
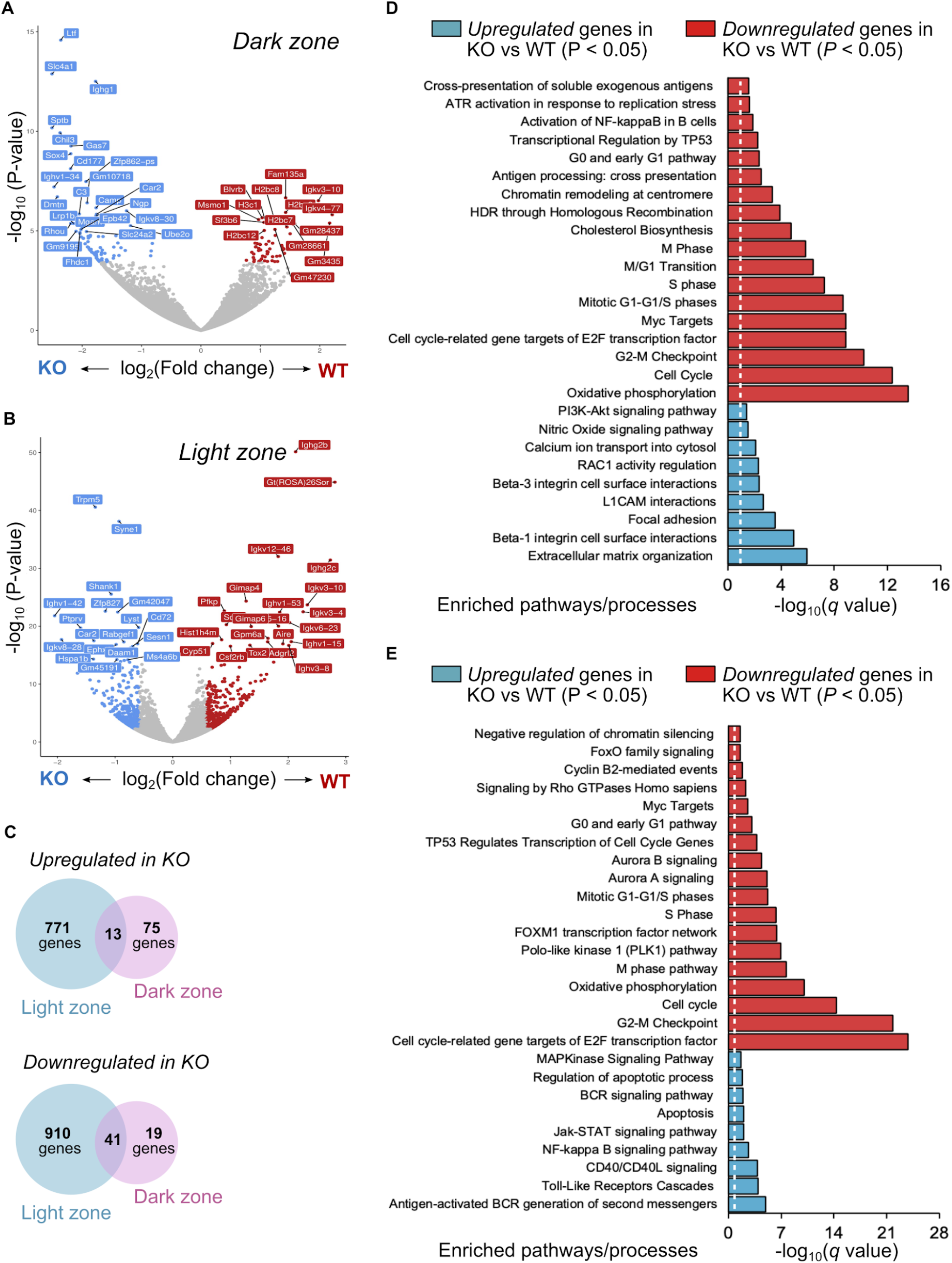
Transcriptional network reveals pathways dysregulated in CD22KO GC B-cells. (**A**) Volcano plot of dysregulated genes in DZ CD22KO GC B-cells. (**B**) Volcano plot of upregulated (blue) and downregulated genes in LZ CD22KO GC B-cells. (**C**) Venn diagram showing the number of significantly dysregulated genes (BH corrected *P* value < 0.05) in CD22KO GC B-cells. (**D**) Gene set enrichment analysis of upregulated (blue) and downregulated genes in DZ CD22KO GC B-cells. (**E**) Gene set enrichment analysis of upregulated (blue) and downregulated genes in LZ CD22KO GC B-cells.

### CD22^KO^ and CMAH^ON^ GC B-cells display hyperactivated BCR signaling

Previous studies demonstrated that GC B-cells display a highly inhibited phenotype when compared with non-GC B-cell populations (*12*, *13*). To interrogate whether this phenotype is partly controlled by CD22, we used an *in vivo* reporter for BCR activation, Nur77-GFP (*39*), to probe the degree of activation between WT and CD22^KO^ GC B-cells. We immunized Nur77^GFP^ (WT control) and CD22^KO^×Nur77^GFP^ mice with OVA displaying liposomes (**Fig. 6A**). After immunization, we found that CD22^KO^ mice produced more GFP^+^ DZ (**Fig. 6B-D**) and LZ (**Fig. 6E-G**) GC B-cells than WT. GFP expression in CD22^KO^ GC B-cells is also evidently higher than WT, suggesting increased activity of the Nur77 promoter in GC B-cells. Consistent with this observation, transcript analysis confirmed increased expression of *Nr4a1*, which encodes for Nur77 (**Fig. 6H**).

We next examined changes in intracellular Ca^2+^ mobilization in CMAH^ON^ or CD22^KO^ GC B-cells in competition with WT GC B-cells, following BCR crosslinking (**Fig. 6I**). Following stimulation with goat F(ab’)_2_ anti-mouse Igκ, we found that CMAH^ON^ GC B-cells (B220^+^CD38^low^PNA^+^) displayed an approximately 2-fold higher Ca^2+^ mobilization than their WT counterparts (**Fig. 6J, K**). In the same tube, naive B-cells (B220^+^CD38^+^PNA^−^) showed no difference between the two genotypes (**Fig. 6L**). Similarly, CD22^KO^ GC B-cells exhibited increased Ca^2+^ flux compared to WT GC B-cells, which appeared to be even more noticeably enhanced than in the naive compartment (**Fig. 6M-O**). Collectively, these findings indicate that CD22 and natural downregulation of its ligands on GC B-cells control the activation of BCR-mediated signaling following antigen engagement.

**Fig 6.**
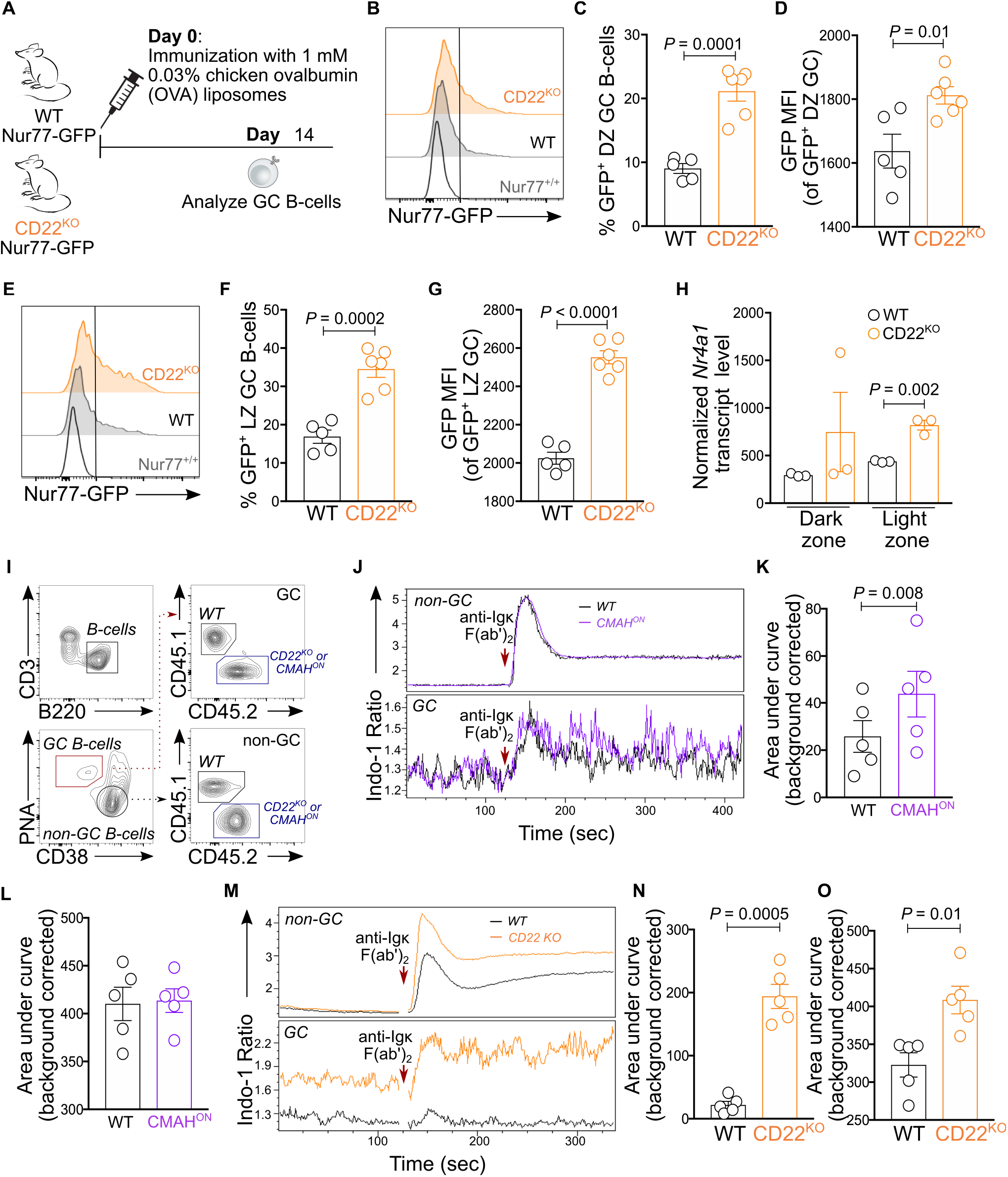
Loss of CD22 or sustained expression of CMAH hyperactivates B-cell activation in the GC. (**A**) Scheme for immunization of WT Nur77-GFP and CD22KO Nur77-GFP mice. (**B-D**) Representative flow cytometric histogram of GFP expression in DZ GC compartment (B) and quantifications of percent GFP+ B-cells in WT and CD22KO DZ GC compartment (C) as well as the GFP MFI of GFP+ populations in WT and CD22KO DZ GC B-cells (D). (**E-G**) Representative flow cytometric histogram of GFP expression in LZ GC compartment (E) and quantifications of percent GFP+ B-cells in WT and CD22KO LZ GC compartment (F) as well as the GFP MFI of GFP+ populations in WT and CD22KO LZ GC B-cells (G). (**H**) Normalized transcript levels of *Nr4a1* gene in DZ and LZ compartments of WT and CD22KO GC B-cells. (**I**) Flow cytometric gating strategy for measurement of intracellular Ca2+ mobilization in GC B-cells. (**J-L**) Time-dependent analysis of Ca2+ flux between WT and CMAHON B-cells using the Indo-1 AM dye (J). Area under the curve (AUC) is plotted for GC (**K**) and non-GC (**L**) compartments. (**M-O**) Time-dependent analysis of Ca2+ flux between WT and CD22KO B-cells using the Indo-1 AM dye (M). Area under the curve (AUC) is plotted for GC (**N**) and non-GC (**O**) compartments.

### GC B-cells from CD22^KO^ and CMAH^ON^ mice display reduced MHCII expression and defective antigen processing

Interaction of GC B-cells with CD4^+^ GC T_FH_ cells is a limiting factor in GC B-cell selection (*40*). To investigate whether CD22 plays a role in the ability of GC B-cell to interact with T_FH_ cells, we first assessed whether surface expression of MHC-II is regulated by CD22 or its *cis*-ligands. Expression of MHC-II in CMAH^ON^ and CD22^KO^ GC B-cells on day 7 post-immunization was comparable with their WT counterparts, however, on day 14 post-immunization, both CMAH^ON^ and CD22^KO^ expressed less surface MHC-II (**Fig. 7A, B**). Interestingly, the transcript levels of the gene encoding for MHC-II (*H2-Ab1*) were comparable between WT and CD22^KO^ GC B-cells, isolated from mice after day 14 post-immunization (**Table S1**). We next checked if this phenotype is due to defective antigen internalization or processing. To interrogate antigen internalization, we assessed the levels of surface BCR following BCR crosslinking with soluble antigen and observed that both CMAH^ON^ and CD22^KO^ GC B-cells displayed similar rates of BCR internalization compared to WT GC B-cells, indicating that CD22 or its *cis*-ligands is dispensable for proper internalization of antigen-BCR complex (**Fig. 7C-F**). Additionally, we explored whether processing of endocytosed antigen is defective in CMAH^ON^ or CD22^KO^ GC B-cells. In order to do this, an *ex vivo* antigen degradation assay was used, which relies on a dsDNA-based sensor that is fluorescently quenched in its undegraded state (*41*). Upon antigen degradation in the lysosome, the dsDNA sensor is degraded, leading to an increase in Atto647 fluorescence. Using this method, we found that both CMAH^ON^ and CD22^KO^ GC B-cells showed significantly lower antigen degradation than their WT counterpart after BCR stimulation (**Fig. 7G-I**). Furthermore, reduced antigen degradation in CMAH^ON^ GC B-cells is dependent on the expression of CD22 on these cells (**Fig. 7J**). Collectively, these findings suggest that loss of CD22 expression or the inability to downregulate CD22 ligands in GC B-cells leads to downregulation of cell surface expression of MHC-II, which may be partly due to impaired antigen processing.

**Fig 7.**
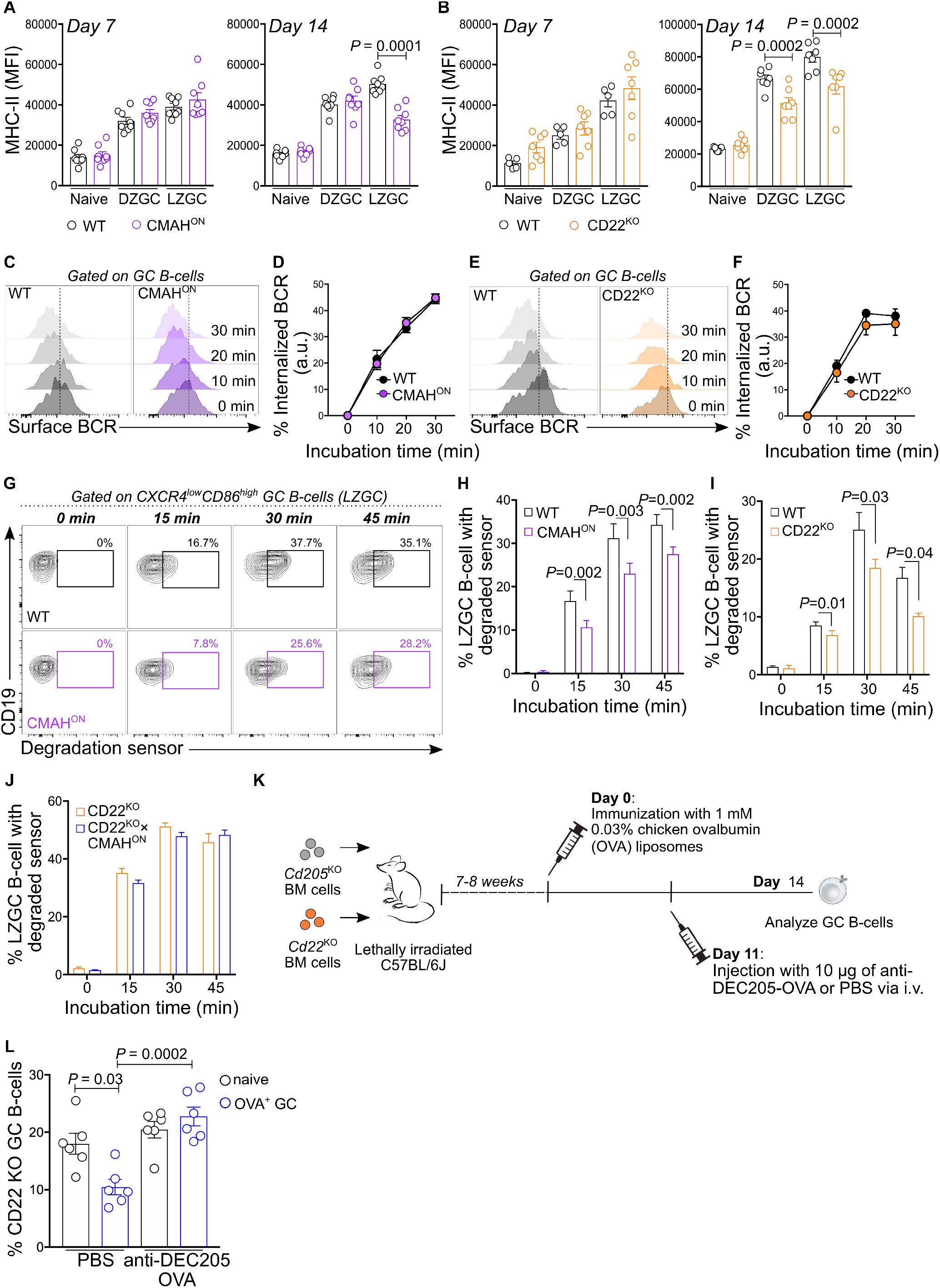
CD22-deficient and CMAH-expressing GC B-cells show reduced surface expression of MHC-II, normal BCR internalization, and lower capacity to degrade internalized antigens. (**A**) Expression of MHC-II on WT and CMAHON naive, DZ, and LZ GC B-cells on days 7 and 14 post-immunization. (**B**) Expression of MHC-II on WT and CD22KO naive and GC B-cells on days 7 and 14 post-immunization. (**C,E**) Representative flow histograms of surface BCR of GC B-cells from immunized mixed chimera mice reconstituted with WT and CMAHON (C) and WT and CD22KO (E) mixed bone marrow cells, at different time points following BCR stimulation with goat F(ab’)_2_ anti-mouse Igκ-biotin. (**D,F**) Quantification of BCR internalization rate from BCR-stimulated WT and CMAHON (D) and WT and CD22KO (F) GC B-cells. (**G**) Representative flow plots tracking the degradation of sensor (increase Atto647 fluorescence) in WT and CMAHON LZ GC B-cells at different time points post-BCR cross linking and incubation at 37oC. (**H-J**) Quantification of degraded sensor in WT and CMAHON (H) WT and CD22KO (I), and CD22KO and CD22KO×CMAHON (J) LZ GC B-cells. (**K**) Scheme used for immunization, and administration of anti-DEC205-OVA or PBS in mixed BM chimera mice reconstituted with BM cells from *Cd205*KO and *Cd22*KO donor mice. (**L**) Quantification of the ratios of CD22KO naive and OVA+ GC B-cells in chimera mice, three days after intravenous injection of either PBS (control) or anti-DEC205-OVA (day 14 post-immunization).

### CD22 does not play a direct role in GC B-TFH interactions

The defective antigen-processing and MHC II expression on CD22^KO^ or CMAH^ON^ GC B-cells suggests that B-T cell interactions may be compromised. To override defective antigen-presentation, we used a previously developed approach to deliver antigen to GC B-cells through a non-BCR-dependent pathway (*42*). Antigen delivered through the DEC205 receptor is known to increase pMHC-II presentation and CD4^+^ T_FH_ interactions. To do so, mixed chimera mice reconstituted with *Cd205*^KO^ (*Cd205* gene encodes for DEC205) and *Cd22*^KO^ BM cells were injected with anti-DEC205-OVA or PBS 11 days after immunization with OVA-displaying liposomes (**Fig. 7K**). Three days later (day 14), we found that administration of anti-DEC205-OVA, but not PBS, restored the levels of antigen-specific CD22^KO^ GC B-cells to a comparable ratio of CD22^KO^ B-cells in the naive population (**Fig. 7L**). These data suggest that CD22 in GC B-cells does not directly impact B-T interactions, but rather plays a critical role in optimal presentation of antigen in a B-cell intrinsic manner.

### CD22 is critical for the proliferation and survival of B-cells in the GC

Robust proliferation and resistance to apoptosis of positively selected B-cells are pivotal in ensuring the maintenance of clones with high-affinity BCRs, and ultimately the production of high-affinity antibodies. To test whether CD22 is important in GC B-cell proliferation, we injected immunized mixed chimeras transplanted with an equal ratio of WT and *Cd22*^KO^ BM cells with BrdU, which labels proliferating cells. Using flow cytometry, we identified a substantially lower number of BrdU^+^ CD22^KO^ GC B-cells than their WT counterparts (**Fig. 8A, B**). However, we did not observe this difference in CMAH^ON^ GC B-cells (**Fig. 8C, D**). We further investigated this difference in proliferation between WT and CD22^KO^ by examining whether it correlates with impaired cell cycle progression, by determining the cell cycle phases using BrdU and 7-AAD (**Fig. 8E**). We found a higher number of CD22^KO^ GC B-cells in the G0-G1 phase (BrdU^−^7-AAD^low^), and a lower number in the S phase (BrdU^+^) and G2-M (BrdU^−^7-AAD^high^) phases compared to WT GC B-cells (**Fig. 8F**). These data are consistent with a defective cell cycle transition in CD22^KO^ B-cells observed from the transcriptome analysis above. Interestingly, we observed a modestly higher number of CMAH^ON^ GC B-cells in both the S and G2-M phases than their WT counterparts (**Fig. 8G**). Together, these findings indicate that CD22 is required for effective expansion of B-cells in the GC.

**Fig 8.**
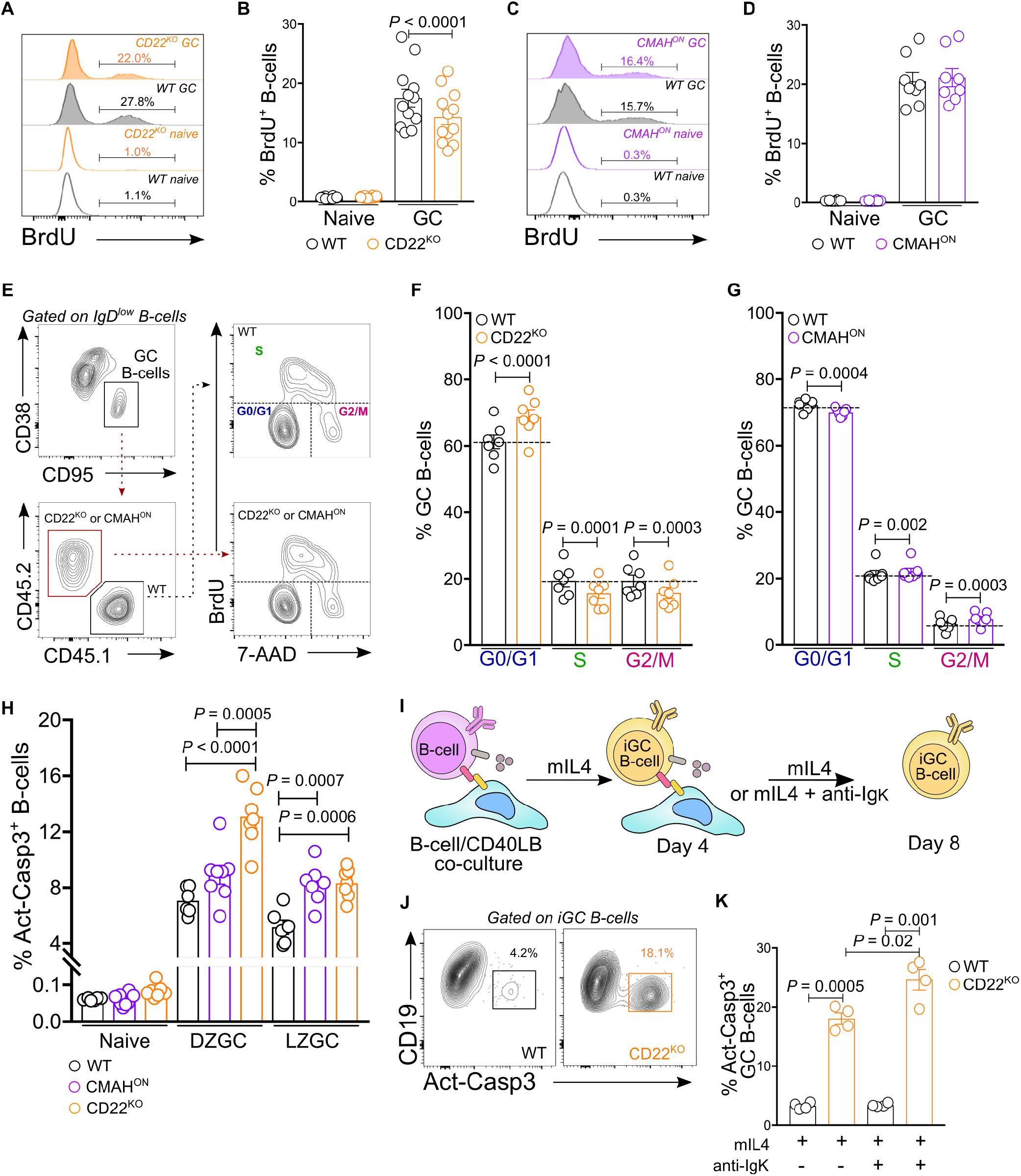
CD22-deficient GC B-cells display poor proliferation and cell cycle phase progression, as well as increased apoptotic cell death. (**A,B**) Representative flow cytometric histograms of BrdU+ cells in WT and CD22KO naive and GC B-cells (A) and quantification of percent BrdU+ cells in WT and CD22KO B-cells (B). (**C,D**) Representative flow cytometric histograms of BrdU+ cells in WT and CMAHON naive and GC B-cells (C) and quantification of BrdU+ cells in WT and CMAHON B-cells (D). (**E**) Gating strategy for the flow cytometric-based analysis of cell-cycle phase progression. (**F,G**) Quantification of WT and CD22KO GC B-cells (F) and WT CMAHON GC B-cells (G) in different cell cycle phases. (**H**) Quantification of percent apoptotic naive, DZ GC, and LZ GC B-cells from WT, CD22KO, and CMAHON mice, at day 14 post-immunization. (**I**) Graphical representation of *in vitro* induced GC B-cell formation and culture. (**J,K**) Flow cytometric gating strategy of apoptotic *in vitro* induced GC B-cells (J) and quantification of percent apoptotic *in vitro* induced WT and CD22KO GC B-cells cultured with or without 1 μg/mL of goat F(ab’)_2_ anti-mouse Igκ.

To investigate whether CD22 or loss of its ligands is necessary for survival of B-cells in the GC, we stained splenocytes from immunized WT, CMAH^ON^, and CD22^KO^ mice, at day 14 post-immunization, with anti-Act-Casp3 antibody, which is an established marker for cells undergoing apoptotic cell death (*43*). We found a higher number of CD22^KO^ B-cells undergoing apoptosis in both the DZ and LZ compartments than WT (**Fig. 8H**). On the other hand, in CMAH^ON^ B-cells, only the B-cells in the LZ compartment displayed heightened apoptosis when compared with WT.

We were interested in understanding whether this increased apoptosis in CD22 is related to hyperactivation of the BCR following stimulation. To examine this, we co-cultured WT or CD22^KO^ naive B-cells with an engineered mouse fibroblast expressing CD40L and BAFF (i.e. CD40LB feeder line) (*44*) in the presence of mIL4 to induce the differentiation of naive (B220^+^CD19^+^IgD^+^) B-cells into GC-like (B220^+^CD19^+^IgD^−^PNA^+^Fas^+^) cells, denoted as iGC. We continued culturing these iGC cells either with mIL4 or mIL4 and anti-mouse Igκ (**Fig. 8I**). Following culture, we found that iGC cells that lack CD22 produced more apoptotic cells in the presence of BCR stimulant (**Fig. 8J, K**). In contrast, WT iGC showed no difference in the levels of apoptotic cells in either culture conditions. This *in vitro* experiment supports the idea that effective dampening of B-cell activation by CD22 is crucial for the survival of B-cells in the GC.

We next tested whether this enhanced apoptosis in CD22^KO^ iGC *in vitro* leads to reduction in induced plasmablast (iPB; CD138^high^CD19^+^ cells) formation. To do this, a 1:1 ratio of WT and CD22^KO^ or CMAH^ON^ naive B-cells were co-cultured onto the CD40LB feeder line. Following establishment of iGC cells in culture, we continued co-culturing cells onto CD40LB cells in the presence of either mIL21 or mIL21 and anti-mouse Igκ. Supplementation of mIL21 induces the differentiation of iGC into plasmablasts *in vitro* (**Fig. S4A-E**) (*44*). After day 8 of co-culture, we found that addition of BCR stimulant in culture resulted in more WT iGC cells differentiating into iPB cells compared with the ratio of differentiated WT iPBs in culture supplemented with mIL21 alone (**Fig. S4F,G**). This finding suggests that enhanced BCR signaling, mediated by loss of CD22 or constitutive expression of CMAH, is detrimental to the differentiation of GC B-cells into plasmablasts *in vitro*.

## Discussion

Affinity maturation of antibodies towards T-dependent antigens relies on the selection of GC B-cells with high affinity BCRs, followed by differentiation into plasma cells. Several lines of evidence have demonstrated that positive selection of B-cells in the GC necessitates optimal signals from BCR and CD40-CD40L interaction to fine-tune expression of c-Myc, a transcriptional regulator critical for cellular proliferation, GC selection, and DZ re-entry (*11*, *45, 46*). GC B-cells strongly dampen the activation signals delivered via the BCR as compared to their naive counterparts, and increasing BCR signal strength is deleterious (*10*, *11*, *41*). Dampened BCR signaling is mediated by recruitment of SHP1 to the BCR complex (*10*, *20*, *21*). However, what enables SHP1/BCR co-colocalization has remained incompletely understood.

Glycans are complex carbohydrate structures covalently bound to protein and lipid carriers. The abundance, size, and composition of these molecules have been correlated to phenotype, activation state, and differentiation of immune cells (*47*–*49*). In this study, we probed the physiological role of glycan remodeling on GC B-cells, which results in the generation of the GL7 epitope, α2-6-linked Neu5Ac, and downregulation of the preferred glycan ligands of CD22 on GC B-cells, which are α2-6-linked Neu5Gc (*24*). Our *in vivo* experiments identified that this specific change in glycosylation is crucial for the maintenance of antigen-specific GC B-cells, egress of plasma cells and memory B-cells, and affinity maturation of antibodies. We postulate that downregulation of CD22 glycan ligands on glycoproteins, such as CD45 and CD22 (*50*–*52*), disrupts the formation and maintenance of CD22 nanoclusters on GC B-cells, leading to a more rapid dissemination of CD22 molecules and attenuation of BCR signaling (**Fig. S5**). Earlier studies investigating the role of glycan ligands on CD22 function on naive B-cells support this hypothesis. Specifically, loss of CD22-ligand interactions was found to enhance colocalization of CD22 and the BCR complex, effectively dampening BCR signaling following antigen engagement (*25*–*27*). Indeed, our *in vivo* experiments confirmed that the defective GC in mice with B-cells constitutively expressing α2-6-linked Neu5Gc is dependent on CD22, highlighting that coordinated downregulation of CD22 ligands acts to fine-tune the BCR signal in the GC by modulating the CD22. Depletion of CD22 on B-cells also showed defective GC and antibody responses despite normal SHM and ability to generate affinity enhancing mutations in the immunoglobulin gene. We further determined that this critical role played by CD22 in the GC is dependent on its ITIMs, which get phosphorylated and recruiting SHP1. Not surprisingly, the GC defects that we observed in CD22^KO^ B-cells are not fully recapitulated in CMAH^ON^ B-cells. In fact, CMAH^ON^ GC B-cells displayed normal proliferation and modest, yet significant, defects in antibody affinity maturation, Ca^2+^ mobilization, and apoptosis when compared to CD22^KO^ GC B-cells. Given that the presence of high affinity ligands for CD22 does not completely inhibit CD22 mobilization, as previously described (*53*), we believe that constitutive expression of high affinity ligands on CMAH^ON^ GC B-cells only partially prevents CD22 interaction with the BCR, resulting in less profound defects than the CD22^KO^ GC B-cells. Nevertheless, our findings support the model in which dampened BCR signaling favours selection and maintenance of GC B-cell clones with high affinity BCRs. Additionally, our results delineate that intrinsic glycan remodeling, leading to loss of high affinity CD22 ligands, during GC B-cell differentiation is a crucial mechanism in regulating the CD22-BCR axis in a T-dependent immune response.

A previous study by Chappell *et al.* (*54*) also described a role for CD22 in the GC reaction. The authors tracked the formation of GC B-cells in mice that had been adoptively transferred with either WT or CD22^KO^ B1-8^hi^ B-cells, which express high-affinity BCR for the hapten NP. While they observed a significant reduction of antigen-specific memory B-cells and plasma cells in immunized mice adoptively transferred with CD22^KO^ B1-8^hi^ B-cells, the authors found comparable levels of GC differentiation and normal antibody responses between WT and CD22^KO^ B-cells. Key differences in the study design between the work by Chappell *et al.* and our current work may explain these seemingly conflicting findings. First, different time points used to analyze the GC response. While we already observed a competitive disadvantage for CD22^KO^ GC B-cells in mixed BM chimera mice on day 7 post-immunization, in the absence of the competition with WT GC B-cells, a GC defect in CD22^KO^ mice was only observed beginning at day 14 post-immunization, which is a week later than Chappell *et al.* investigated. The second difference between the two studies is the difference in B-cell models. The B1-8^hi^ B-cells used in the earlier study have a high affinity BCR. It is thought that GC B-cell clones with high affinity BCRs capture more antigens, leading to stronger T-cell help (*55, 56*), which could potentially rescue the detrimental impact of CD22 deletion. On the other hand, the majority of our studies examined the formation of GCs from the endogenous antigen-specific B-cells, which typically express BCRs with low-to-moderate affinities to their cognate antigens.

A recent study by Luo *et al.* (*57*) explored the molecular mechanism downstream of the BCR complex that may be altered, leading to attenuated BCR signaling in GC B-cells. The authors identified that GC B-cells have an altered AKT-mediated signaling network, which was pinpointed to differential AKT T308 and S473 phosphorylation. Proteomic analysis revealed that AKT kinase activity is rewired in GC B-cells to selectively target negative regulators of BCR signaling, such as SHP1, CSK, and HPK1. Our study demonstrated another intrinsic remodeling that occurs in GC B-cells that may be linked to reduced BCR signal activation. By tracking Nur77 promoter activity through GFP reporter expression and *ex vivo* intracellular Ca^2+^ mobilization following BCR crosslinking, we found that CD22 and loss of its *cis*-ligands, in part, maintains a hypoactivated phenotype of GC B-cells. Furthermore, we demonstrated that enhanced BCR signaling in GC B-cells lacking CD22 or constitutively expressing the preferred CD22 ligands is linked to increased GC B-cell apoptosis and reduced plasma cell differentiation. Several factors may be in play to promote activation-induced cell death (AICD) in GC B-cells. A study by Akkaya *et al.* (*55*) demonstrated that BCR activation increased oxidative phosphorylation and glycolytic activities in B-cells. However, prolonged BCR signaling, resulting in intracellular reactive oxygen species (ROS) and Ca^2+^ accumulation, impaired mitochondrial function, leading to AICD. Strikingly, the presence of a second signal coming from cognate T-cells were able to rescue both the mitochondrial dysfunction and AICD. Findings from our transcriptome analysis revealed that besides controlling the BCR activation, CD22 is also essential for proper metabolic function and expression of many mitochondrial enzymes. Thus, we speculate that enhanced BCR signal in CD22^KO^ and CMAH^ON^ GC B-cells may have induced a defective mitochondrial function; and for GC B-cells with low-to-moderate affinity BCRs, which are believed to have a competitive disadvantage for GC TFH help, this dysfunction may have resulted in their eventual demise.

Strength of GC B-cell interaction with their cognate T_FH_ cells dictates the differentiation of GC B-cells into plasma cells (*9, 58*). As it was suggested that B-T interaction is enhanced in a CD22-dependent manner (*59*), we investigated whether CD22 plays a pivotal role in B-T interaction in the GC. We determined that the surface expression of MHC-II is substantially reduced in CD22^KO^ and CMAH^ON^ GC B-cells. Forcing GC B-T_FH_ interaction *in vivo*, by administration of anti-DEC-antigen following the establishment of the GC, rescued the disadvantage of not having CD22, suggesting that CD22 does not play a direct role in B-T interaction. However, given that we only compared the GC response of CD22^KO^ B-cells following PBS or anti-DEC205-OVA treatment with a negative control DEC205^KO^ B-cells, the experiment does not fully conclude that loss of CD22 does not impair GC B-T_FH_ interaction. This warrants further investigations. One way to address this is by comparing the GC response between WT and CD22^KO^ B-cells after anti-DEC205-OVA administration. This assay might be able to precisely detect any modest impairment in T_FH_ interaction, if there is any. Another way would be to interrogate whether CD22 is involved in CD40L surface mobilization or proper secretion of GC-relevant cytokines.

Maintenance of positively selected GC B-cells mainly depends on its ability to cycle between the LZ and DZ and sustaining a robust and fast proliferation while in the DZ. Following GC T_FH_ interaction, GC B-cells activate the mechanistic target of rapamycin complex 1 (mTORC1) and cMyc transcription factor to support cell growth and cell cycle progression of B-cells in the GC (*45, 46, 60*). Our transcriptome analysis uncovered that CD22 is required by GC B-cells to properly regulate the cell cycle machinery, which includes E2F regulated genes, Aurora and PLK1 kinase signaling, and cell cycle checkpoint genes. Using *in vivo* BrdU labeling, we found that CD22^KO^ GC B-cells proliferate less and display impaired cell cycle progression, validating the transcriptome finding. While loss of CD22 has been correlated with hyperproliferation in naive B-cells through the activation of the PI3K-AKT pathway, it is possible that constrained BCR signaling, partly controlled by CD22, is responsible for maintaining a rewired AKT activity in the GC. Interestingly, AKT kinase is rewired in the GC B-cells to specifically target proteins relevant to cell cycle progression, such as E2F targets (e.g. PCNA, CBX5, and NASP), G2/M checkpoints (e.g. AURKB, NUP50, and RBM14), and Myc targets (e.g. RACK1, PA2G4, and DEK) (*57*). Thus, we can speculate that enhanced BCR signaling through CD22 deletion may lead to altered AKT activity in the GC and loss of substrate selectivity. Another explanation for the reduced proliferation of CD22^KO^ GC B-cells can be derived from our transcriptome data. DZ CD22^KO^ GC B-cells displayed an impaired cholesterol biosynthesis pathway. Cholesterol is a critical component of the cell membrane and the ability of GC B-cells to synthesize cholesterol *de novo* may have structural and signaling benefits, especially in the DZ compartment where B-cells induce cellular growth and division (*61*–*63*).

In summary, our findings describe a functional role for the emergence of the glycan epitope for the GL7 antibody. We define that appearance of the GL7 epitope is not merely a defining marker for the GC B-cell population, but represents a change in CD22 ligands, which plays an important role in fine-tuning BCR signaling within the GC B-cells. We conclude that loss of CD22 ligands is crucial for the maintenance and selection of GC B-cells, as well as affinity maturation antibodies by fine tuning inhibitory function of CD22 on GC B-cells.

## Materials and Methods

### Mouse strains and frozen bone marrow cells

All mice used in this study were on a C57BL/6J genetic background. C57BL/6J (stock no.:000664), B6-CD45.1 (stock no.:002014), *Mb1^Cre^* (stock no.:020505), and *Aidca^Cre^* (stock no.:007770) mice were acquired from Jackson Laboratory. Hy10 mice were a gift from Dr. Jason Cyster (UC San Francisco, USA). Frozen bone marrow cells from *Cd205*^KO^ mice were provided to us by Dr. Gabriel Victora (Rockefeller University, USA). Bone marrow cells were tested for mouse pathogens (Charles River) prior to use.

*Rosa26^lsl^-Cmah* mice were generated following a previously described protocol (*64–66*). To confirm the genotype of *Rosa26^lsl^-Cmah* mice, PCR was performed using extracted DNA from ear notch samples. The WT Rosa26 locus was identified using 5’-GGAGCGGGAGAAATGGATATG-3’ as forward primer and 5’-AAAGTCGCTCTGAGTTGTTAT-3’ as reverse primer (band size: 600 bp). On the other hand, the *Rosa26^lsl^-Cmah* gene was determined using 5’-ATTCTAGTTGTGGTTTGTCC-3’ and 5’-ATGGTGCTCACGTCTAACTTCC-3’ primers (band size: 370 bp).

Floxed *Cd22* mice *(C57BL / 6N-Cd22tm1a(EUCOMM)Wtsi/Tcp)* were rederived from sperm acquired through The Centre for Phenogenomics (Toronto, Canada). Initially, mice hemizygous for the tm1a allele were established and crossed with Flp deleter mice (B6.129S4-*Gt(ROSA)26Sortm2(FLP*)Sor*/J) from The Jackson Laboratory. Successful conversion of the tm1a allele into the tm1c allele of mouse *Cd22*, which we denote as *Cd22*^f/f^, was verified by PCR for the presence of a proximal *loxP* site (primers: 5’-AAACAGCATGGGCTCTGCTTCACAG-3’ and 5’-TGGAGAGGCAAGCAAAGATGGAGAGG-3’; band size: 102 bp) and distal *loxP* site (primers: 5’-GCGCAACGCAATTAATGATAAC-3’ and 5’-TACAGTCATTTGAAAGAGGCCAGC-3’; band size: 211 bp). The WT *Cd22* locus was identified using 5’-GCGGGAAGGGACTGGCTGCTAT-3’ and 5’-AGTCCAGAGACCATCGGCAAG-3’ primers (band size: 270 bp), which also recognized the tm1c allele. Mice were subsequently backcrossed onto C57BL/6J mice to remove the flp recombinase, and crossed with the relevant strains expressing Cre to generate mice homozygous for *Cd22^f/f^* and bearing either *Mb1^Cre^* or *Aidca^Cre^*.

All mice used in this study were bred and maintained under specific pathogen-free conditions. Experiments described in the study were approved by the Health Sciences Animal Care and Use Committee of the University of Alberta.

### Bone marrow chimera and adoptive transfer

To generate mouse chimeras, WT recipients were irradiated with 1000 rad and injected intravenously with isolated bone marrow cells from various donor mice within 24 hours. Mixed bone marrow chimera mice were generated by reconstituting lethally irradiated WT mice with 1:1 ratio, unless otherwise specified, of bone marrow cells from indicated donor mice. Chimera mice were used after 7-10 weeks following bone marrow transplantation. For adoptive transfer of HEL-specific B-cells, B-cells from spleens of Hy10 mice were isolated using a mouse B-cell negative-isolation kit (Miltenyi). Approximately 1×10^6^ splenic B-cells were injected into B6 recipient mice intravenously a day before immunization.

### Cell lines

Human embryonic kidney (HEK) 293T cells were acquired from ATCC and cultured in DMEM media supplemented with 10% fetal bovine serum (FBS; Gibco) and 100 U/mL of penicillin and 100 μg/mL streptomycin (Gibco). Chinese hamster ovary (CHO) Flp-In cells (Invitrogen) were maintained in DMEM-F12 media supplemented with 5% FBS, 100 U/mL of penicillin/100 μg/mL streptomycin, and 10 mM HEPES. The CD40LB feeder cell line was a gift from Dr. Daisuke Kitamura (Tokyo University of Science, Japan) and Dr. Javier Marcelo Di Noia (University of Montreal, Canada). Feeder cells were cultured in DMEM supplemented with 10% FBS and 100 U/mL of penicillin/100 μg/mL streptomycin.

### Liposome preparation, immunization, and tamoxifen administration

Lyophilized lipids containing a 65:35:5 molar ratio of distearoyl phosphatidylcholine (DSPC) (Avanti Polar Lipids), cholesterol (Sigma), and PEG_2000_-distearoyl phosphoethanolamine (PEG_2000_-DSPE; Avanti Polar Lipids) were hydrated in sterile PBS and sonicated for a minimum of 20 minutes. NP-OVA-PEG_2000_-DPSE, OVA-PEG_2000_-DPSE or HEL-PEG_2000_-DPSE, which were generated using a previously published protocol (*67*), was added to the lipid mixture at the time of hydration. The molar fraction of Protein-PEG_2000_-DPSE were maintained at 0.03% in all liposome preparations. Following sonication, liposomes were passed at least 15 times through 800-nm and 100-nm filters using a hand-held extrusion device (Avanti Polar Lipids). Excess Protein-PEG_2000_-DPSE was separated from liposomes using a CL-4B Sepharose (Sigma) column. Eluted liposomes were diluted in sterile PBS to 1 mM concentration prior to administration to mice.

For immunization involving liposomes, each mouse received 200 μL of 1 mM liposomes via tail vein injection. Where indicated, at day 11 post-immunization with OVA-liposomes, mice were boosted with 10 μg of anti-DEC205-OVA, prepared according to a published study (*42*), via tail vein injection. Spleens were collected at different time points post-immunization for microscopy and flow cytometric analyses of the GC.

For temporal expression of human CD20-driven Cre recombinase, mouse chimeras reconstituted with *CD20-ERT2^Cre^* and *CD20-ERT2^Cre^*×*R26^lsl^-Cmah* BM cells were gavaged with 1mg tamoxifen (Combi-Blocks) in 200 μL corn oil (Sigma) on days 8 and 9 post-immunization with OVA displaying liposomes. Spleens were collected on Day 14 (5 days post-tamoxifen) for flow cytometric analysis of GC B-cells.

### Flow cytometry and sorting

Spleens were macerated in RPMI 1640 containing 2% FBS and passed through a 40 μm mesh. Single cell suspensions were treated with Ammonium-Chloride-Potassium (ACK) lysis buffer for five minutes at 4°C to deplete red blood cells (RBC). RBC-depleted splenocytes were washed and resuspended in HBSS containing 2% FBS or 0.1% bovine serum albumin (flow buffer). Aliquots of cell suspensions were stained on ice for 30 minutes using appropriate reagents and surface marker antibodies as listed in **Table S3**. Stained cells were washed and resuspended in flow buffer supplemented with 1 μg/mL of viability dye propidium iodide or 7-AAD. Naive and GC B-cell compartments were gated on a Fortessa X-20 flow cytometer as B220^+^CD19^+^IgD^+^ and B220^+^CD19^+^IgD^−^ CD38^low^Fas^+^GL7^+^(or PNA^+^) live single cells, respectively, unless specified otherwise. OVA-specific GC B-cells were further gated on the flow cytometer using OVA-AF647 (Invitrogen). NP^+^ GC B-cells were determined in flow using NP_20_-APC and NP_4_-APC (prepared according to protocol provided by Dr. Pierre Milpied of CIML, France). Dark zone and light zone compartments were identified as CXCR4^high^CD86^low^ and CXCR4^low^CD86^high^ GC B-cells, respectively.

For intracellular staining, cells stained with surface marker antibodies were fixed and permeabilized with BD Cytofix/Cytoperm™ (BD Biosciences) on ice for 20 minutes. Fixed cells were washed two times with flow buffer, followed by incubation with intracellular marker antibodies for 30 minutes at 4°C.

For cell cycle and proliferation assays, immunized mice were injected intravenously with 1 mg BrdU (Sigma) dissolved in sterile PBS. Spleens were harvested after 1 hour and RBC-depleted single cell suspensions were stained with surface marker antibodies for naive and GC B-cells for 30 minutes at 4°C. Following incubation, cells were washed with flow buffer and fixed with BD Cytofix/Cytoperm™ for 20 minutes at 4°C. Cells were then washed 2 times with flow buffer and incubated with BD Cytoperm™ Permeabilization Buffer Plus (BD Biosciences) for 10 minutes at 4°C. Cells were washed twice with flow buffer, re-fixed with BD Cytofix/Cytoperm™ for 10 minutes at 4°C, and incubated at 37°C with 20 μg/mL of DNase I (Sigma) in PBS containing Ca^2+^ and Mg^2+^. Following incubation, cells were washed and stained with anti-BrdU antibody (BioLegend) and 7-AAD for 20 minutes at room temperature. Proliferating cells were gated on the flow cytometer as BrdU^+^ B-cells. Cell cycle phases of GC B-cells were identified as described previously (*68*).

For sorting GC B-cells, RBC-depleted splenocytes were stained with appropriate surface marker antibodies in HBSS containing 2% of FBS and 1mM EDTA (FACS buffer) for 30 minutes at 4°C. Cells were washed and resuspended in FACS buffer supplemented with 1 μg/mL 7-AAD. GC B-cells were sorted by BD FACS Aria III (BD Biosciences) using a 100-μm nozzle. Sorted single cells were then collected in cold RPMI 1640 media containing 10% FBS, 5.5×10^−5^ M 2-mercaptoethanol (Sigma), 10mM HEPES (Gibco), 1mM sodium pyruvate (Gibco), 1mM essential amino acids (Gibco), 100 U/mL of penicillin and 100 μg/mL streptomycin.

### Immunofluorescence

Spleens from non-immunized and immunized mice were embedded in cryomatrix (Thermo Fisher), froze, and stored at -80°C prior to sectioning. Cryostat tissue sections (9 μm) were fixed and permeabilized in a cold 1:1 mixture of acetone and methanol for 5 minutes, blocked with HBSS supplemented with 5% FBS for 40 minutes at room temperature, and washed with PBS-T. Tissue sections were then stained with lectin PNA-biotin (Vector Lab) and antibodies GL7-biotin (BioLegend), anti-mouse IgD (clone: 11.26c.2a; BioLegend), anti-mouse CD21/35 (clone: 7G6; BD Biosciences) in different combinations overnight at 4°C.The following day, tissue sections were washed 3 times with PBS-T and the following secondary antibodies were used to detect the primary antibodies: Streptavidin-AF555 (Invitrogen), anti-rat IgG2a-AF488 (SouthernBiotech), and anti-rat IgG2b-AF647 (BioLegend). Following incubation, tissue sections were washed 5 times with PBST and stained with 2.5 μg/ml Hoechst for 15 minutes at room temperature. Images of spleen sections were acquired with either Zeiss LSM700 microscope (20× magnification) or Molecular Devices ImageXpress Pico (10× magnification). Individual GC clusters were imaged with Zeiss LSM700 confocal microscope using 63× magnification. Image analysis was performed using ZEN Blue and Fiji processing software.

### Enzyme-linked immunosorbent assay (ELISA)

To measure the affinity maturation of antigen-specific antibodies, sera were collected on days 14, 21, and 42 after immunization with 0.03% NP-OVA liposomal nanoparticles. High affinity NP-specific and total NP-specific antibodies were captured on plates pre-coated with 10 μg/mL of NP_2_-BSA (prepared following the recommended protocol provided by Dr. Daisuke Kitamura) and NP_23_-BSA (LGC Biosearch), respectively. Plates were washed 5 times with PBS-T and incubated with goat anti-mouse IgG1 antibody conjugated with horseradish peroxidase (SouthernBiotech) for 1 hour at room temperature. Plates were then washed 5 times with PBS-T followed by addition of peroxidase substrate 2,2’-azino-bis(3-ethylbenzothiazoline-6-sulfonic acid) (ABTS) solution (SeraCare) into each well for colour development. After 15 minutes, reactions were stopped with 1M phosphoric acid. Optical density was collected using a microplate reader set to 405 nm.

### RNAseq and transcriptome analysis

Total RNA from sorted GC B-cells of immunized WT and CD22^KO^ mice was extracted using an RNeasy micro kit (Qiagen). Indexed cDNA libraries were generated using a SMART-Seq Stranded kit (Takara Bio) for the Illumina platform and the quality of each constructed library was measured using a Bioanalyzer 2100 (Agilent). Paired-end 150 bp sequencing was performed on an Illumina HiSeq platform by Novogene. For transcriptome analysis, the quality of unaligned reads was first validated using FastQC v0.11.2 (*69*) followed by the removal of Illumina adapters using Cutadapt v3.3 (*70*). Adapter-trimmed sequencing reads were mapped to the GRCm38/mm10 reference genome using HISAT2 (*71*). Aligned reads were counted and annotated to Gencode M25 (*72*) using HTseq-count (*73*), followed by read count normalization and differential expression analysis by DESeq2 (74). Differentially expressed genes (DEGs) were determined based on the Benjamini-Hochberg (BH) corrected P value ≤ 0.05. Gene enrichment analysis was performed on DEGs with an unadjusted P value ≤ 0.05 using EnrichR (75–77) and default settings. Significantly enriched gene sets were identified based on BH P value < 0.05.

### Ig VH 186.2 DNA sequencing and mutation analysis

Approximately 30,000 GC B-cells (B220^+^CD19^+^IgD^−^GL7^+^NP_20_-APC^+^ live cells) were isolated using FACS from splenocytes of WT and CD22^KO^ mice, at day 21 after immunization with NP-OVA liposomes. Genomic DNA was extracted using a DNA microprep kit (Qiagen). To sequence the V_H_186.2 region, V_H_186.2 was amplified from the extracted genomic DNA by PCR using 5’-AGCTGTATCATGCTCTTCTTGGCA-3’ and 5’-AGATGGAGGCCAGTGAGGGAC-3’ as forward and reverse primers, respectively. PCR was performed following a previously published (*78*). PCR products were then cloned into a pBAD vector (Invitrogen) and transformed into chemically competent DH5α *E. coli* (NEB Biolabs). Plasmids were extracted from *E. coli* clones using a GeneJET plasmid miniprep kit (Thermo Fisher), and DNA was sequenced using a Sanger sequencer. Mutational analysis of the Ig V region was performed using the IMGT/V-QUEST tool (https://www.imgt.org).

### Anti-DEC205-OVA plasmids and recombinant protein expression and purification

Plasmids encoding the anti-DEC205 light chain and anti-DEC205 heavy chain fused to OVA were transformed into chemically competent DH5α *E. coli* and purified using a GeneJET plasmid miniprep kit (Thermo Fisher). To express anti-DEC205-OVA, HEK293T cells were cultured in complete DMEM in T175 flasks and incubated at 37°C in a 5% CO_2_ incubator until they reached approximately 70% confluence. HEK293T cells in each flask were then added with 2 mL of OptiPro SFM containing 10 μg each of anti-DEC205 light chain and heavy chain plasmids, and 40 μg of branched polyethylenimine (Sigma) and incubated overnight at 37°C. The following day, the media were replaced with fresh complete DMEM and cultured for another 48-72 hours. Media were collected, filter sterilized, and stored at 4°C prior to purification. Anti-DEC205-OVA was purified from the media using a HiTrap Protein G HP column (GE Healthcare) according to the manufacturer’s recommendation, dialyzed in PBS, and quantified as described previously (*42*).

### Generation of mouse CD22-Fc fusion

To generate the mouse CD22-Fc fusion, DNA encoding 1-435 amino acid residues of mouse CD22 was amplified by PCR using 5’-AGCAGCGCTAGCATGCGCGTCCATTACCTGTGG-3’ and 5’-AGCAGCACCGGTATCCAGCTTAGCTTCCTG-3’ as forward and reverse primers, respectively. PCR products were cloned upstream of the human IgG1 Fc domain encoded in the pcDNA5/FRT/V5-His-TOPO^®^ vector, which was described elsewhere (*79*). Sequence verified plasmid encoding for mCD22-Fc was transfected together with Flp recombinase-expressing plasmid pOG44 (Invitrogen) using lipofectamine LTX (Invitrogen) into chinese hamster ovary (CHO) Flp-In cells. Stably transfected cells were selected after 10 days of culturing in DMEM-F12 containing 5% FBS, 100 U/mL of penicillin/100 μg/mL streptomycin, 2.438 g/L sodium bicarbonate (Gibco) and 1 mg/mL Hygromycin B (Invitrogen). For expression of mCD22-Fc, approximately 1×10^6^ of mCD22-Fc expressing CHO cells were plated onto a T175 flask in DMEM-F12 media supplemented with 5% FBS, 100 U/mL of penicillin/100 μg/mL streptomycin, and 10 mM HEPES. Cells were then incubated at 37°C for 10-12 days. Supernatant from cultured cells was collected and filtered using a Rapid-Flow™ sterile filter with 0.2 μm PES membrane pore size (Nalgene).

To generate a non-binding mutant of mouse CD22-Fc (mCD22 R124A-Fc), site-directed mutagenesis was performed on the sialic acid-binding region of CD22 using a megaprimer-PCR strategy (*80*). Briefly, the megaprimer containing the R124A mutation was generated by PCR using 5’-AGCAGCGCTAGCATGCGCGTCCATTACCTGTGG-3’ as the forward primer and 5’-AGTCCCTGCGGTCATGGCCAACCCCAGATTCCC-3’ as the reverse primer. A second round of PCR was performed to amplify the 1-435 amino acid residues of mouse CD22 containing the R124A mutation, using the megaprimer and 5’-AGCAGCACCGGTATCCAGCTTAGCTTCCTG-3’ as the forward and reverse primers, respectively. Cloning and protein expression of mCD22 R124A-Fc follow the same protocol as mentioned above.

### In vitro GC B-cell culture

For induction of GC B-cells *in vitro*, B-cells were isolated from mouse spleens using FACS or a mouse B-cell isolation kit (Miltenyi) and co-cultured with CD40LB fibroblast feeder lines in RPMI 1640 medium supplemented with 10% FBS, 5.5 × 10^−5^ M 2-mercaptoethanol, 10mM HEPES, 1mM sodium pyruvate, 1mM essential amino acids, 100 U/mL of penicillin, and 100 μg/mL streptomycin. Mouse IL-4 (1 ng/mL; BioLegend) was added to primary co-culture for 4 days. Induced GC B-cells were then replated onto a fresh CD40LB feeder line and cultured with mouse IL-21 (10 ng/mL; BioLegend) or mouse IL-4 for another 3 to 4 days.

### Intracellular Ca^2+^ mobilization

Approximately 15×10^6^ splenocytes from mice, at day 14 after immunization, were resuspended in HBSS (Gibco) supplemented with 1% FBS and 1 μM Indo-1 AM (Invitrogen), and incubated at 37°C for 30 minutes. Cells were washed with a 5-fold volume of HBSS with 1% FBS and centrifuge at 300 rcf for 5 minutes. Pelleted cells were stained with surface marker lectin PNA and antibodies against mouse B220, CD38, CD45.1, and CD45.2 for 20 minutes at 4°C. Cells were washed and resuspended in HBSS containing 1% FBS. A 0.5 mL aliquot containing approximately 1 × 10^6^ splenocytes was incubated for 5 minutes at 37°C prior to the Ca^2+^ mobilization assay. Ca^2+^ mobilization in GC and non-GC B-cells populations was detected using a Fortessa X-20 flow cytometer. After baseline was established (approximately 2-3 minutes), splenocytes were stimulated with buffer (1% FBS/HBSS) or 10 μg/mL goat F(ab’)_2_ anti-mouse Igκ (SouthernBiotech) and Indo-1 fluorescence (violet vs blue) was monitored by flow cytometry for 3-5 minutes at 37°C. Data were analyzed using the kinetics function on FlowJo v 9.

### Ex vivo antigen internalization and processing assay

For the BCR internalization assay, RBC-depleted splenocytes were incubated with 1 μg/mL of goat F(ab’)_2_ anti-mouse Igκ-biotin (SouthernBiotech) for 30 minutes at 4°C. Splenocytes were washed 3 times with cold HBSS containing 0.1% BSA and resuspended in pre-warmed (37°C) RPMI 1640 medium supplemented with 10% FBS, 5.5 × 10^−5^ M 2-mercaptoethanol, 10mM HEPES, 1mM sodium pyruvate, 1mM essential amino acids, 100 U/mL of penicillin, and 100 μg/mL streptomycin. Cells were then incubated in a 5% CO_2_ incubator at 37°C for specified timepoints. At the end of each timepoint, cells were immediately fixed in 1% paraformaldehyde (PFA) for 10 minutes at room temperature. Fixed cells were washed 3 times with 0.1% BSA in HBSS followed by incubation of surface marker antibodies and fluorochrome-conjugated streptavidin. Changes in the levels of anti-mouse Igκ-biotin on fixed naive and GC B-cells were monitored by flow cytometer.

For the antigen degradation assay, splenocytes from immunized mice were stained with 1 μg/mL of goat F(ab’)_2_ anti-mouse Igκ-biotin followed by 1 μg/mL of streptavidin (BioLegend) at 4°C. Cells were washed 3 times with HBSS containing 0.1% BSA and incubated with 100 nM of biotin-conjugated antigen degradation sensor described in Nowosad et al. (*41*) at 4°C for 20 minutes. Stained cells were then washed 3 times, resuspended in pre-warmed RPMI 1640 medium supplemented with 10% FBS, 5.5 × 10^−5^ M 2-mercaptoethanol, 10mM HEPES, 1mM sodium pyruvate, 1mM essential amino acids, 100 U/mL of penicillin, and 100 μg/mL streptomycin, and incubated in a 5% CO2 incubator at 37°C for specified timepoints. After each timepoint, cells were fixed and stained with GC B-cell relevant surface marker antibodies. Changes in the fluorescence of the antigen degradation sensor were measured by flow cytometry

### Statistical analysis

All statistical analyses in this study were done with GraphPad Prism software except for the calculation of statistical significance of DEGs and enriched pathways, which were both performed in DESeq2 (*74*) and EnrichR (*75*–*77*), respectively. For experiments comparing two groups, a two-tailed Student’s *t*-test was used to evaluate statistical significance. When the assay was carried out in competition, a paired Student’s *t*-test was used; while an unpaired Student’s *t*-test was performed for samples analyzed independently. For experiments with more than 2 sample groups, a one-way analysis of variance (ANOVA) was performed to assess differences between means. Tukey or Dunnet multiple comparisons tests were used to determine statistical significance between groups.

## Data and code availability

Next generation sequencing data are deposited in the Gene Expression Omnibus (GEO) database. No new codes were generated for the transcriptome analysis.

## Acknowledgements

We thank Dr. Daisuke Kitamura (Tokyo University of Science, Japan) and Dr. Javier Marcelo Di Noia (University of Montreal) for providing the CD40L feeder cell lines; Dr. Juhee Pae and Dr. Gabriel Victora (Rockefeller University, USA) for providing anti-DEC205-OVA plasmids and frozen bone marrow cells from *Cd205*^KO^ mice; Dr. Jason Cyster (UC San Francisco, USA) for access to Hy10 mice; and Sudip Subedi of the Applied Genomics Core at the University of Alberta for processing RNA for next generation sequencing analysis. We thank Pierre Milpied for critical reading and feedback of our manuscript. We would also like to thank the University of Alberta’s Flow Cytometry Core for sorter use and Health Sciences Laboratory Animal Services for animal husbandry and technical assistance.

## Author contributions

J.R.E. and M.S.M. designed the experiments, analyzed and interpreted the data, and wrote the manuscript. J.R.E. performed all mouse and *ex vivo* experiments. S.S. performed mouse genotyping and assisted with all aspects of the mouse experiments. L.S. carried out experiments with *in vitro* induced GC B-cells. J.J. cloned and expressed mCD22-Fc. B.M.A. and J.C.P. helped in developing the *R26^lsl^-Cmah* and *Cd22^f/f^* mice, and in interpreting preliminary data from these models. S.J.M. and L.N. provided bone marrow cells from knock-in mice containing ITIM-deficient CD22. C.X. and H.T. provided reagents and expert advice for establishing *R26^lsl^-Cmah* mice. T.A.B. provided Nur77^GFP^ mice and advised on their use and interpretation of the data. M.J.S. provided the *CD20-ERT2^Cre^* mice and provided critical advice on the direction of the project. All authors have read and approved the manuscript.

