## Supplementary Information for "Coordinated changes in glycosylation regulate the germinal centre through CD22"

### Supplementary Materials

#### Supplementary Tables

**Table S1. Deseq2 normalized count of genes in mouse dark zone and light zone WT and CD22 KO GC B-cells (spreadsheet).**

**Table S2. Differentially expressed genes between WT and CD22 KO GC B-cells (spreadsheet).**

**Table S3. List of antibodies, recombinant proteins, chemicals, and other resources used in this study.**

| Reagent or resource | Source | Identifier |
|---|---|---|
| **Antibodies** | | |
| Rat anti-mouse B220 (clone:RA3-6B2) | BD Biosciences | cat# 563793 |
| Rat anti-mouse CD19 (clone: 6D5) | BioLegend | cat# 115555 |
| Rat anti-mouse CD38 (clone: 90) | BioLegend | cat# 102732, 102716, 102722 |
| Hamster anti-mouse CD19 (clone: Jo2) | BD Biosciences | cat# 563646 |
| Rat anti-mouse IgD (clone: 11-26c.2a) | BD Biosciences | cat# 563618 |
| Rat anti-mouse IgD (clone: 11-26c.2a) | BioLegend | cat# 405718 |
| Rat anti-mouse CD86 (clone: GL1) | BD Biosciences | cat# 560582 |
| Rat anti-mouse/human GL7 antigen (clone: GL7) | BioLegend | cat# 144606, 144610, 144620 |
| Rat anti-mouse CD184 (CXCR4) (clone: L276F12) | BioLegend | cat# 146506, 146511, 146517 |

| | | |
|---|---|---|
| Mouse anti-mouse CD45.1 (clone: A20) | BioLegend | cat# 110738, 110736, 110718 |
| Mouse anti-mouse CD45.2 (clone: 104) | BioLegend | cat# 109816, 109832, 109841, 109836 |
| Rat anti-mouse I-A/I-E (clone: M5/114.15.2) | BioLegend | cat# 107639 |
| Mouse anti-BrdU (clone: 3D4) | BioLegend | cat# 364108 |
| Rat anti-mouse CD138 (clone: 281-2) | BD Biosciences | cat# 561070, 563147 |
| Rat anti-mouse CD22 (clone: OX-97) | BioLegend | cat# 126112, 126108, 126106 |
| Mouse anti-mouse CD22.2 (clone: Cy34.1) | BD Biosciences | cat# 740682 |
| Goat anti-mouse IgG1-HRP | SouthernBiotech | cat# 1071-05 |
| Goat F(ab')$_2$ anti-mouse Kappa UNLB | SouthernBiotech | cat# 1052-01 |
| Goat F(ab')$_2$ anti-mouse Kappa Biotin | SouthernBiotech | cat# 1052-08 |
| Rat anti-mouse CD21/CD35 (clone: 7G6) | BD Biosciences | cat# 553817 |
| Mouse anti-rat IgG2b (clone: MRG2b-85) | BioLegend | cat# 408209 |
| Mouse anti-mouse CD45.1 (clone: A20) | BioLegend | cat# 110738, 110736, 110718 |
| Mouse anti-rat IgG2a (clone: 2A8F4) | SouthernBiotech | cat# 3065-30 |
| Chicken anti-Neu5Gc | BioLegend | cat# 146903 |

| | | |
|---|---|---|
| Goat anti-chicken IgY secondary antibody | Invitrogen | cat# A21447 |
| **Chemicals and recombinant proteins** | | |
| FITC-Peanut Agglutinin (PNA) | Vector Laboratories | cat# FL-1071 |
| Biotin-PNA | Vector Laboratories | cat# B-1075 |
| AF647-Chicken ovalbumin (OVA) | Invitrogen | cat# O34784 |
| Purified Streptavidin | BioLegend | cat# 280302 |
| AF488-Streptavidin | BioLegend | cat# 405235 |
| Brilliant violet 605-streptavidin | BioLegend | cat# 405229 |
| Brilliant violet 650-streptavidin | BioLegend | cat# 405232 |
| Propidium iodide | Invitrogen | cat# P1304MP |
| 7-AAD | BioLegend | cat# 420403 |
| OVA | Sigma | cat# A-5503 |
| $NP_{23}$-BSA | LGC Biosearch | cat# N-5050H |
| $NP_{2}$-BSA | Prepared in the lab | N/A |
| $NP_{4}$-APC | Prepared in the lab | N/A |
| $NP_{20}$-APC | Prepared in the lab | N/A |
| NP-OVA | Prepared in the lab | N/A |
| NP-OSu | LGC Biosearch | cat# N-1010-100 |

| | | |
|---|---|---|
| Tamoxifen | Combi-Blocks | cat# QC-0156 |
| 1,2-distearoyl-sn-glycero-3-phosphocholine (DSPC) | Avanti Polar Lipids | cat# 850365P |
| 1,2-distearoyl-sn-glycero-3-phosphoethanolamine-N-[methoxy(polyethylene glycol)-2000] (ammonium salt) ($PEG_{2000}$-DSPE) | Avanti Polar Lipids | cat# 880120 |
| Cholesterol | Sigma | cat# C8667 |
| 4% paraformaldehyde | Thermo Scientific | cat# J19943-K2 |
| Polyethylenimine, branched | Sigma | cat# 408727 |
| Lysozyme, chicken egg white (HEL) | Sigma | cat# 62971 |
| N-succinimidyl 3-(2-pyridyldithio)-propionate (SPDP) | Thermo Scientific | cat# 21857 |
| Maleimide-$PEG_{2000}$-DSPE | Avanti Polar Lipids | cat# 880126 |
| Peroxidase substrate solution A | SeraCare | cat# 50-76-02 |
| Peroxidase substrate solution B | SeraCare | cat# 50-65-02 |
| Sepharose CL-4B | GE Healthcare | cat# 17-0150-01 |
| Mouse IL4 | BioLegend | cat# 574304 |
| Mouse IL21 | BioLegend | cat# 574504 |
| **Critical buffers, kits, and other resources** | | |
| Mouse B-cell isolation kit | Miltenyi | cat# 130-090-862 |

| | | |
|---|---|---|
| LD columns | Miltenyi | cat# 130-042-901 |
| CytoFix/CytoPerm buffer | BD Biosciences | cat# 554714 |
| CytoPerm buffer plus | BD Biosciences | cat# 51-2356KC |
| HiTrap Protein G HP column | GE Healthcare | cat# 17-0404-01 |
| CountBright™ absolute counting beads | Invitrogen | cat# 36950 |
| Fiji | ImageJ | https://imagej.net/Fiji |
| FlowJo v9 | FlowJo | https://www.flowjo.com |
| GraphPad Prism v7 | GraphPad Software | https://www.graphpad.com/scientific-software/prism/ |
| ZEN blue | Zeiss | https://www.zeiss.com/ |
| Galaxy platform (RNAseq analysis) | Galaxy | https://usegalaxy.org/ |

### Supplementary Figures

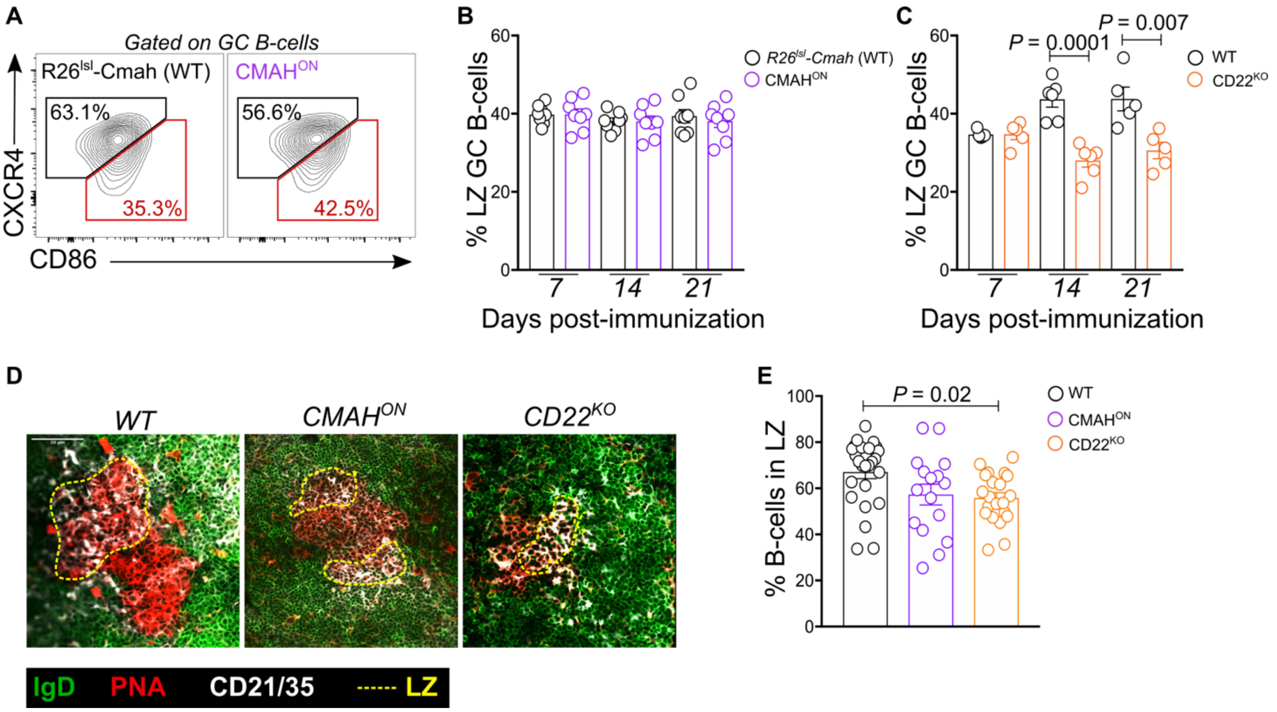


**Fig. S1. Loss of CD22 impairs DZ-to-LZ transition of GC B-cells**. (**A**) Flow cytometric gating strategy for identification of DZ and LZ GC B-cells. (**B**) Quantification of percent LZ B-cells in *R26$^{lsl}$-Cmah* (WT) and CMAH$^{ON}$ GCs at varied timepoints post-immunization. (**C**) Quantification of percent LZ B-cells in WT and CD22$^{KO}$ GCs at varied timepoints post-immunization. (**D,E**) Immunofluorescence images of LZ compartment (CD21/35$^{+}$) of individual GC clusters in spleens of immunized WT, CMAH$^{ON}$, and CD22 KO mice, at day 14 post-immunization (D) and quantification of % B-cells in each GC cluster (E).

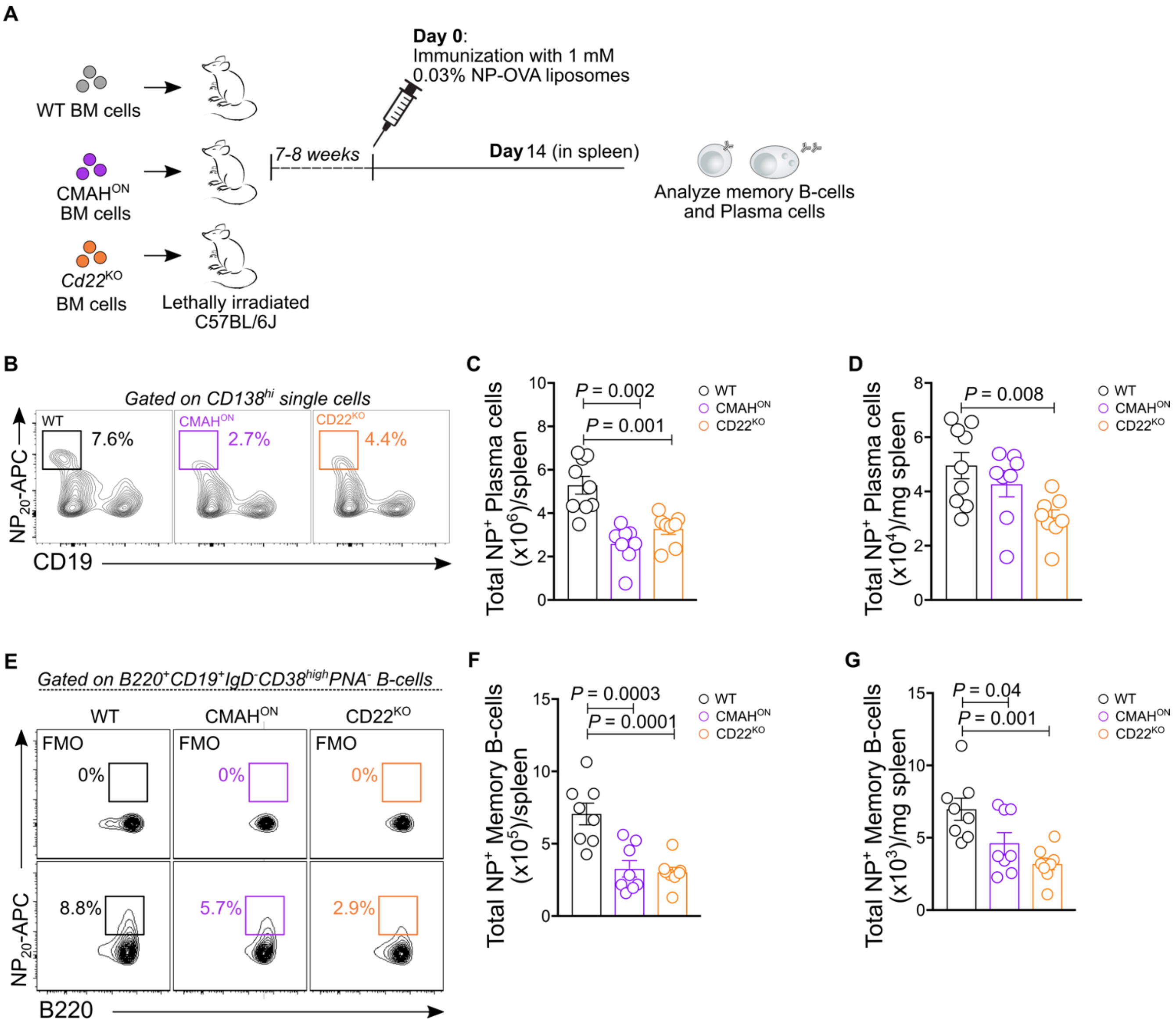


**Fig. S2. CD22$^{KO}$ and CMAH$^{ON}$ GC B-cells display reduced antigen-specific plasma cells or memory B-cells following immunization.** (**A**) Schematic for chimera mice generation and immunization. (**B**) Representative flow plot for gating NP$^{+}$ Plasma cells from fixed and permeabilized splenocytes. (**C,D**) Quantification of total NP$^{+}$ Plasma cells per spleen (C) or mg of spleen (D) at day 14 after immunization. (**E**) Representative flow plots for gating NP$^{+}$ memory B-cells from splenocytes of immunized chimera mice. (**F,G**) Quantification of total NP$^{+}$ memory B-cells per spleen (F) or mg of spleen (G) at day 14 after immunization.

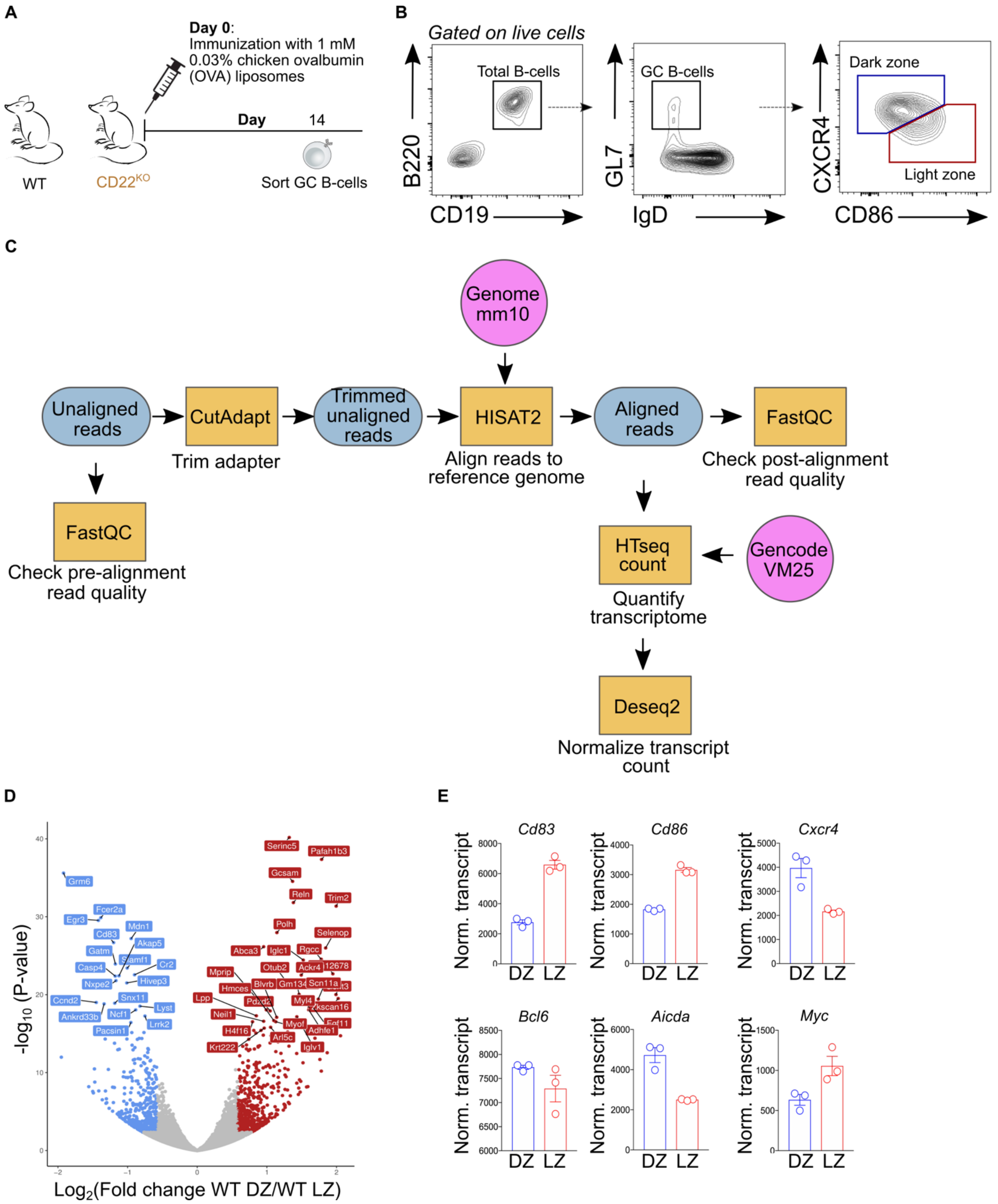


**Fig. S3. Gating strategy for RNAseq analysis.** (**A**) Schematic for immunization. WT and CD22$^{KO}$ mice were immunized with OVA-displaying liposomes intravenously. Spleens were collected on day 14 post-immunization for GC B-cell sorting. (**B**) Gating strategy used in this study to sort DZ and LZ GC B-cells. (**C**) Schematic of reference-based RNAseq analysis used in this study. (**D**) Volcano plot of differentially expressed genes in WT DZ and WT LZ GC B-cells. (**E**) Expression of select GC relevant genes in WT DZ and WT LZ GC B-cells.

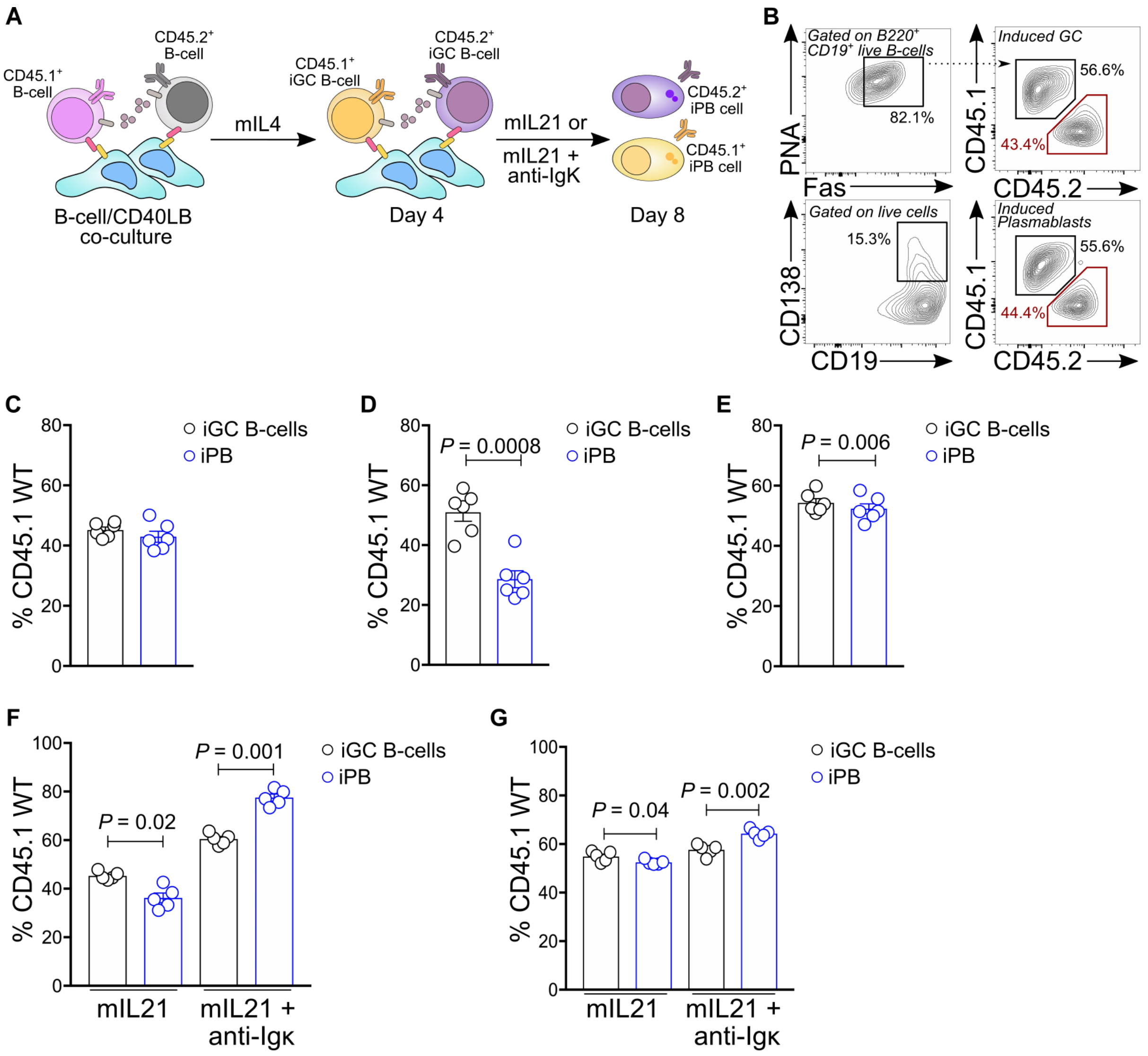


**Fig. S4. BCR signaling impairs plasmablast differentiation of CD22 KO and CMAH-expressing GC B-cells in an *in vitro* co-culture system.** (**A**) Schematic of *in vitro* co-culture of isolated B-cells with CD40LB feeder line. A 1:1 ratio of CD45.1$^{+}$ WT and CD45.2$^{+}$ WT or CD22$^{KO}$ or CMAH$^{ON}$ naive B-cells were plated onto the CD40LB feeder line. Mouse IL4 were added in the media on the first 4 days to induce the formation of GC-like (iGC) B-cells. On day 4, Induced GC B-cells were replated onto a fresh CD40LB feeder line and cultured for another 4 days in media containing either mouse IL21 or mouse IL21 and anti-Igκ to induce iGC differentiation into plasmablasts (iPB) cells. (**B**) Flow gating strategy used to identify *in vitro* induced GC B-cells and plasmablasts cells after day 8 of co-culture. (**C-E**) Quantification of percent CD45.1 WT GC B-cells and plasmablasts that differentiated in co-cultures, which initially contained 1:1 of CD45.1 WT and CD45.2 WT naive B-cells (C), CD45.1 WT vs CD22$^{KO}$ naive B-cells (D), and CD45.1 WT and CMAH$^{ON}$ naive B-cells (E). (**F,G**) Quantification of percent CD45.1 WT GC B-cells and plasmablasts in culture initially containing 1:1 CD45.1$^{+}$ WT and CD22$^{KO}$ (E) and 1:1 CD45.1 WT naive B-cells and CMAH$^{ON}$ naive B-cells (F), and cultured in media containing either mouse IL21 or mouse IL21 and 1 μg/mL of anti-mouse Igκ.

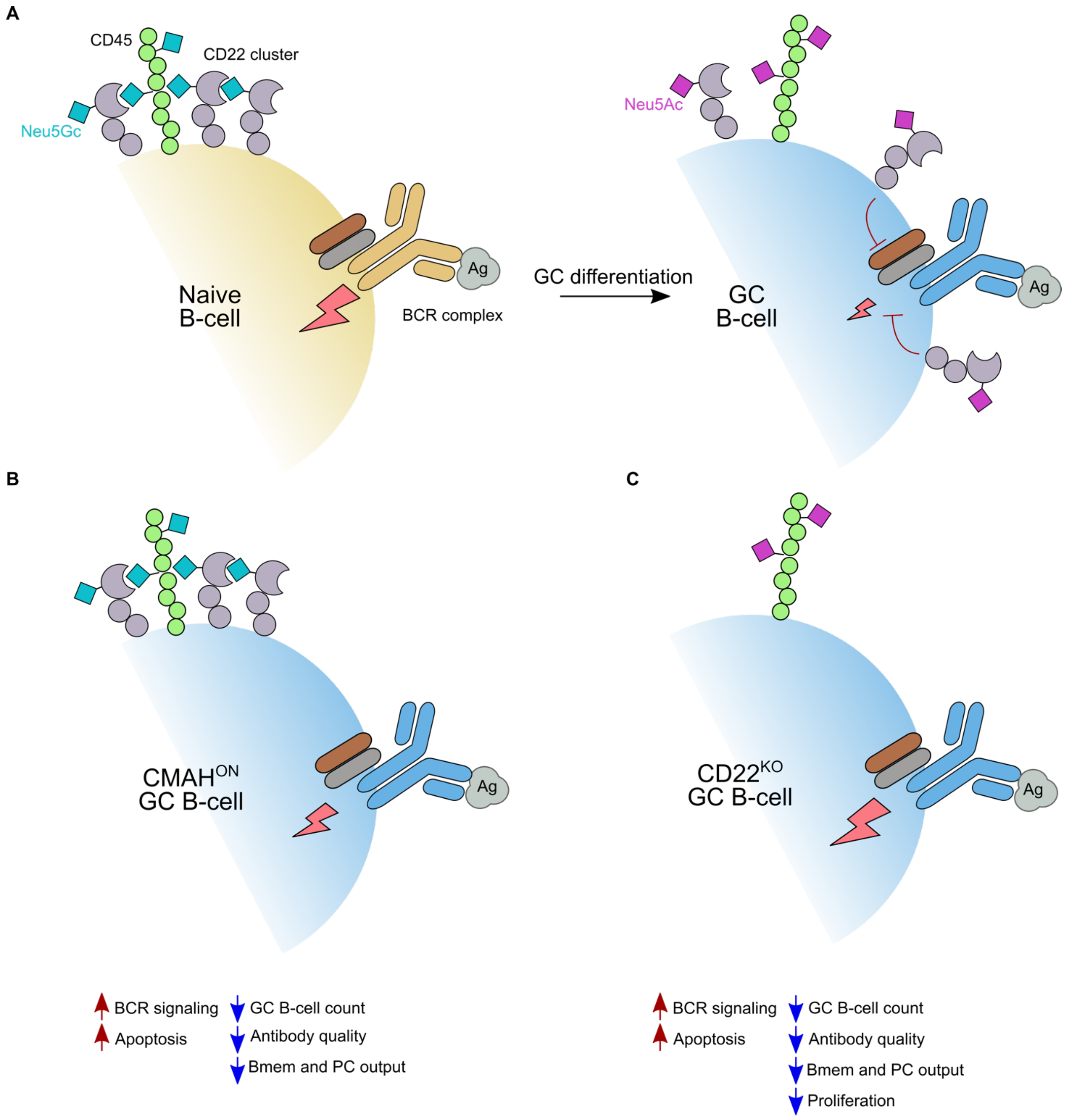


**Fig. S5. Model for the role of glycan remodeling on GC B-cells.** (**A**) Remodeling of sialic acid from Neu5Gc to Neu5Ac as B-cell differentiate in the GC. (**B**) Impact of constitutive CMAH expression in GC B-cell response. (**C**) Impact of CD22 deficiency in GC B-cell response.
